# Polymer Model Integrates Super-Resolution Imaging and Epigenomic Sequencing to Elucidate the Role of Epigenetic Reactions in Shaping 4D Chromatin Organization

**DOI:** 10.1101/2024.10.08.617296

**Authors:** Vinayak Vinayak, Ramin Basir, Rosela Golloshi, Joshua Toth, Lucas Sant’Anna, Melike Lakadamyali, Rachel Patton McCord, Vivek B Shenoy

## Abstract

Chromatin, with its complex spatial and temporal organization, plays a crucial role in regulating gene expression. Recent advancements in super-resolution microscopy have revealed that nanoscale domains of heterochromatin (repressed segments) embedded within a euchromatin (active segments) background are fundamental units of 3D chromatin organization. In tissue-resident cells, the size of these heterochromatin domains varies with the microenvironment, particularly its stiffness, and chromatin organization is also influenced by pharmacological and epigenetic drugs. However, the mechanisms governing heterochromatin domain size under various conditions and their impact on gene expression remain unclear. To address this knowledge gap, we have developed a dynamic, next-generation sequencing informed chromatin copolymer model. Our model simulates the spatiotemporal evolution of chromatin, driven by passive diffusion and active epigenetic reactions, which interconvert euchromatin and heterochromatin. By integrating chromatin-chromatin interaction energetics and diffusion-reaction dynamics, we predict the formation of nanoscale heterochromatin-rich domains and establish a scaling relationship between their size and the modulation of epigenetic reaction rates. Additionally, our model predicts that epigenetic and chromatin compaction changes in response to changes in global reaction rates occur predominantly at domain boundaries. We validated these predictions via Hi-C contact map analysis and super-resolution imaging of hyperacetylated melanoma cells. Subsequent RNA-seq analysis suggested a pivotal role of these epigenetic shifts in influencing the metastatic potential of these cells. We further validated our mesoscale findings against chromatin rearrangement in hMSCs, which exhibit sensitivity of epigenetic reaction rates to changes in microenvironmental stiffness. Finally, we evaluated the effects of cycling of epigenetic reaction rates in silico, mimicking the cellular transition to different extracellular conditions, and back again. This finding reveals a cell-type invariant mechanism driven by domain boundaries, whereby chromatin organization guides epigenetic memory formation. Our findings show that chromatin reorganization in response to changes in epigenetic reaction rates resulting from alterations in the microenvironment, drug exposure and disease progression impacts both immediate cellular responses and long-term epigenetic memory.

## 1 Introduction

The three-dimensional organization of the mammalian genome within the nucleus is a critical determinant of cell fate, regulating transcription and thereby influencing development, differentiation, metabolism, and proliferation. The segregation of active and repressed genes into distinct phases, comprising loosely packed, transcriptionally active, euchromatin (A-compartment) and compact, transcriptionally repressed, heterochromatin (B-compartment) phases, has been established by complimentary approaches[5, 6], including next-generation sequencing techniques (Hi-C[4, 7–9], Micro-C[10], ATAC-seq[11]) and super-resolution imaging[12–14]. Epigenetic profiling experiments such as ChIP-seq[15, 16] and immunofluorescence imaging have established that phase-separated regions exhibit unique epigenetic signatures[17, 18]. Specifically, heterochromatin is characterized by persistent histone methylation (such as H3K9me3 or H3K27me3), whereas euchromatin is characterized by prominent histone acetylation (such as H3K27ac)[3, 19–21]. These marks are instrumental in governing transcriptional likelihood[22]. Super-resolution imaging has revealed the widespread presence of heterochromatin-rich packing domains as fundamental units of chromatin organization[1, 23, 24]. These domains have been observed across diverse cell lines, although this aspect of chromatin organization remains poorly understood.

Chromatin organization exhibits dynamic responsiveness to a broad spectrum of physical and chemical extracellular cues, orchestrating a complex interplay between epigenetic remodeling and transcriptional regulation. Recent studies have elucidated the pivotal role of mechanical stimuli, including substrate elasticity[1] and viscosity[25], in directing the redistribution of histone epigenetic marks via the action of histone epigenetic remodelers such as histone deacetylases (HDACs) and histone methyltransferases (HMTs) (e.g., HDAC3 and EZH2), which are also known as “histone writers” and “histone erasers” [1, 25–28]. Furthermore, diseased states such as fatty liver and cancer metastasis, characterized by altered extracellular lipid concentrations[29] and intra/extravasation[30], also exhibit large scale chromatin reorganization. Notably, pharmacological, and epigenetic interventions evoke similar genomic and epigenomic changes[1], underscoring the central role of epigenetic remodelers in modulating histone marks through methylation and acetylation. These epigenetic changes, in turn, reconfigure nanoscale chromatin domains[31–34], influencing transcriptional outcomes and cell fate. Despite significant advances, the physical mechanisms underlying this multiscale process remain poorly understood, warranting further investigation to elucidate the intricate relationships among chromatin dynamics, epigenetic regulation, and disease pathogenesis.

Modeling efforts in recent years have largely viewed chromatin organization as a passive, equilibrium system, yielding valuable insights into the interplay between epigenetic marks and 3D chromatin structure [35–41]. However, a smaller subset of studies has explored chromatin organization in a more realistic setting, incorporating the active energy flux driven by epigenetic remodelers, which better captures the dynamic nature of chromatin organization. Computational models have begun to explore such active dynamics[42–45], including the impact of cohesin-mediated chromatin looping[46–50] and transcriptional activity[51–53] on chromatin organization, including the phenomenon of epigenetic spreading[45, 54, 55]. However, there is a significant gap in the understanding of how epigenetic reactions, which are influenced by changes in the microenvironment, drug exposure, disease progression, and aging, regulate 4D chromatin organization. This omission overlooks the crucial role of histone epigenetic remodelers in shaping chromatin organization through the formation of heterochromatin-rich packing domains and the role of such an organization in controlling the cell’s transcriptional activity, underscoring the need for a more comprehensive biophysical approach to capture this complex interplay.

Here, we present a chromatin copolymer model that integrates bioinformatics inputs with epigenetic reaction-driven dynamics to predict spatio-temporal genomic organization. The polymer comprising euchromatin and heterochromatin segments is informed by experimentally obtained Hi-C or ChIP-seq data, ensuring a biologically realistic representation and facilitating direct comparison with experimental observations. Our model captures polymer energetics through chromatin-chromatin interactions, and passive dynamics through a combination of nucleoplasmic and epigenetic diffusion, driving phase separation. Epigenetic reactions capture the effect of histone epigenetic remodelers, actively interconverting euchromatin and heterochromatin. This reaction-driven approach enables us to capture the effects of both mechanical and drug-induced changes on chromatin organization. We demonstrate that a delicate balance between the diffusion of epigenetic marks and epigenetic reactions gives rise to characteristic chromatin domains and establish a scaling relationship between their size distribution and reaction rates. We validate these findings through super-resolution (STORM) imaging of chromatin domains in human A375 malignant melanoma cells after hyperacetylation through an epigenetic drug, which reveals the global decompaction of chromatin domains. Bulk Hi-C sequencing of these cells shows excellent agreement with our model predictions, capturing epigenetic compartmental shifts with high accuracy at a kilobase-scale resolution. Notably, our model predicts that epigenetic changes primarily occur at heterochromatin domain boundaries, which is validated by Hi-C contact maps. Further RNA-seq analysis revealed a correlation between chromatin decompaction and the differential regulation of EMT-related genes in melanoma, near domain boundaries. To validate our model’s predictive power in diverse microenvironments, we simulated changes in chromatin organization in human mesenchymal stem cells in response to altered substrate stiffness and experimentally confirm these predictions via super-resolution imaging. Finally, our chromatin polymer model reveals the role of chromatin organization in the formation of history-dependent epigenetic memory, providing a general framework for understanding how extracellular cues shape temporal chromatin organization and gene expression.

## 2 Methods

### 2a Data-informed polymer model of chromatin

Chromatin exhibits complex organization across multiple scales, as revealed through conformation capture maps such as Hi-C and Micro-C[56]. In particular, the A (euchromatin-like and active) and B (heterochromatin-like and repressed) compartments emerge as prominent structures characterized by enhanced chromatin interactions within their respective types. These interactions are physically facilitated by bridging proteins such as HP1[57] and PRC1[58] in addition to other physical interactions, which lead to large-scale phase separation[59, 60]. Simultaneously, experimental investigations such as histone ChIP-seq[61] have demonstrated a significant overlap between active histone tail marks (e.g., H3K27ac) and the A compartment, whereas repressive histone tail marks (e.g., H3K9me3 or H3K27me3) tend to associate with the B compartments[4]. Based on these findings and other computational studies[36, 42], we choose to represent chromatin as a block copolymer, with active/A and repressed/B regions. Specifically, we represent chromatin as a self-avoiding beads-on-a-string polymer, where the beads can be labeled as active/euchromatin-like (blue) or repressed/heterochromatin-like (red) (Fig. 1a). Each bead in the simulation corresponds to a continuous genomic region of 10 kilobase pairs (kb), with a bead size (σ) set at 65 nm. More details on the polymer configuration are provided in SI1.1-1.2.

**Figure 1:**
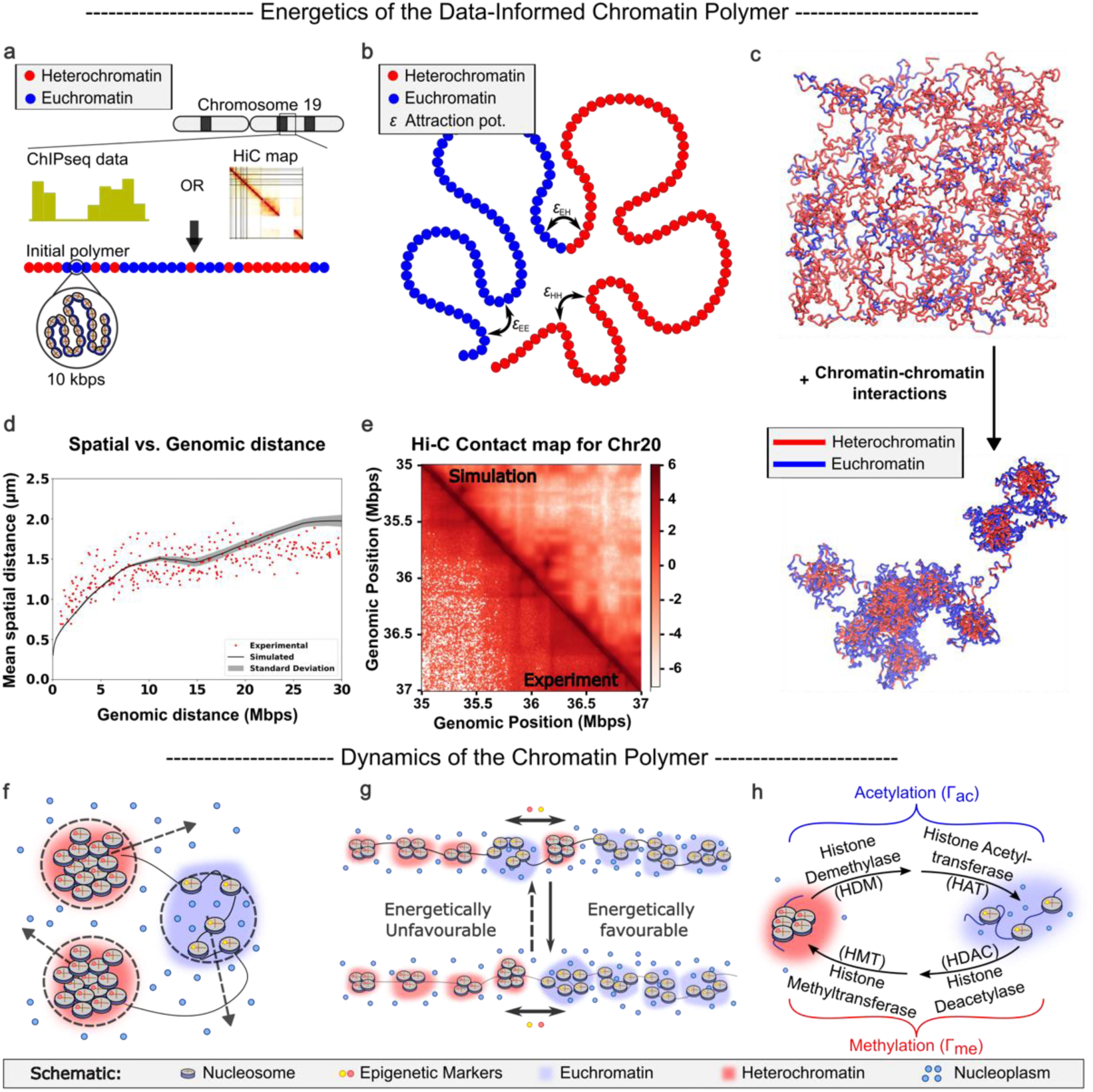
Polymer model based on experimentally obtained inputs and biologically relevant dynamics. a) The initial polymer strand is constructed using inputs from experimentally obtained sequencing data. The histone ChIP-seq data (through ChromHMM[2]) or Hi-C data areused to label the initial configuration of the polymer. We assign all repressed states to heterochromatin (red) and active states to euchromatin (blue) beads. b) Lennard‒Jonespairwise potentials are defined for each pair of beads basedontheir epigenetic marking. *∈_EE_*,*∈_EH_ and ∈_HH_* are the euchromatin-euchromatin, euchromatin-heterochromatin and heterochromatin-heterochromatin bead interaction potential strengths,respectively. c) The system is relaxed (bottom) from a random initial configuration(top) viaLangevin dynamics. d) Mean spatial distance vs. mean genomic distance obtained after polymer relaxation is plotted with the experimental data[3]. e) The simulatedHi-C map (upper triangle) shows thecharacteristic checkerboard pattern,which is observedin the experimental Hi-C map (lower triangle)[4]. f) Diffusion of thenucleoplasm (water) is accounted for implicitly viaBrownian dynamics. g) Diffusion of epigenetic marks is modeled viaan energy-based metropolis criterion for exchanging epigenetic marks between spatially neighboring beads. h) Epigenetic reactions of acetylation (through the action of histone demethylase(HDM) and histone acetyltransferase(HAT)) and methylation (through the action of histone deacetylase(HDAC) and histone methyltransferase(HMT)) are modeled as MonteCarlo-based epigenetic reassignment processes.

We adopt a two-pronged approach to study the genome multiscale organization. First, we consider a prototype polymer of 3000 randomly marked (60% repressed) beads. This polymer is not biased based on any biological input and is used to elucidate the key biophysical principles that govern the formation and evolution of heterochromatin-rich packing domains. Next, to make experimentally testable predictions, we employ a data-driven strategy to label the initial polymer (Fig. 1a) corresponding to chromosomes 19-22 of hMSC cells, leveraging the available Broad ChromHMM data[62]. For chromosomes 18-21 of A375 melanoma cells, the labeling information is constructed from the A/B compartmentalization obtained from Hi-C contact maps. The two approaches for initializing the polymer, data-agnostic and data-driven, complement each other and help validate the robustness of the proposed model. More details on the labeling of the chromatin segments are provided in SI1.1. While more than two epigenetic flavors derived from different ChromHMM states, multiple histone ChIP-seq datasets or finer Hi-C or MicroC compartmentalization can be incorporated into the model, our focus remains on highlighting the key role of heterochromatin domains in regulating changes in gene expression; hence, we stick to a two-component model.

We deliberately keep the polymer energetics simple with minimal sets of parameters that are easy to interpret. To account for the interactions between the chromatin segments, we adopt a Lennard‒Jones interaction potential function, which captures the influence of the bridging proteins and other nonspecific interactions involved in establishing chromatin-chromatin affinities (Fig. 1b). The depth of the Lennard Jones potential well (*ε*) is fixed for each pair of epigenetic marks. Only excluded volume interactions are considered between differently labeled beads to keep the number of parameters small, which leaves us with *∈*_*HH*_ and *∈*_*EE*_, the intra-epigenetic interactions between repressed (heterochromatin-like) beads and active (euchromatin-like) beads, respectively. For *∈*_*EE*_ and *∈*_*HH*_, we constrain our search such that *∈*_*EE*_ < *∈*_*HH*_, which results in a higher packing density for heterochromatin or repressed chromatin regions[21, 63, 64]. A finite extensible nonlinear elastic (FENE) potential is employed between adjacent beads on the polymer to preserve the integrity of the polymer. More details on the potentials can be found in SI1.2. Following this setup, the polymer is relaxed via Langevin dynamics to establish its initial configuration (Fig. 1c), which is the starting point for our production runs. These energetics are able to replicate the biologically relevant, experimentally observed[3] spatial distance-genomic distance relationship (Fig. 1d) and reproduce the signature checkerboard Hi-C contact pattern (Fig. 1e), which is a trademark of A-B spatial compartmentalization[4]. A crucial aspect that we do not include in the following results is the role of cohesin-dependent loop extrusion, which is bounded by stopping factors in the form of CTCF-binding sites. Our model has the capacity to account for this, but at the current level of coarse graining, we do not expect significant deviations in the obtained results (details in SI1.1).

### 2b Dynamics for the temporal evolution of the chromatin polymer

Having defined the energetics of the chromatin segments, we consider the dynamics of spatial and epigenetic reorganization over time. The spatial reorganization of the chromatin, i.e., the displacement of the chromatin in a background of viscous nucleoplasm, is driven by Brownian diffusion of the polymer beads. Concurrently, the epigenetic mark of the polymer beads can passively redistribute on the chromatin polymer, pushing the system to a lower energy configuration, where beads with similar epigenetic marks prefer to phase separate and coarsen. We term this process the diffusion of epigenetic marks and account for it via a Monte Carlo-based epigenetic remarking algorithm. Finally, active epigenetic remodeling is captured by implementing epigenetic reactions. The details of these processes and their molecular dynamics-based execution are discussed in more detail below:

#### Brownian diffusion

The spatial diffusion of chromatin within the nucleoplasm is captured by employing Brownian dynamics to simulate the diffusion of chromatin beads (Fig. 1f). Considering a viscosity of 150 cp[42, 65] and a bead size of 65 nm, we estimate a Brownian time step *τ*_*Br*_ of 0.3 s (calculations provided in SI1.3). To simulate, we choose an integration time step Δ*t* of 0.01*τ*_*Br*._This choice allows us to simulate ∼1 hour of real-time behavior for every 1 × 10^6^ integration steps.

#### Diffusion of epigenetic marks

Given that interactions between heterochromatic regions are more favorable than those between heterochromatin and euchromatin, it is unfavorable for isolated heterochromatin segments to be in a euchromatin-rich environment. When the overall ratio of the hetero and euchromatic beads remains constant, the system tends toward a state with lower energy, leading to the phase separation of euchromatin and heterochromatin. To illustrate this further, we designed a schematic, as shown in Fig. 1g, where the fully phase-separated configuration is more favorable than the other configurations because of unfavorable interactions between chromatin segments. An energetically driven exchange between neighboring regions occurs as we transition from the top to the bottom configuration. This exchange lays the foundation for the diffusion of epigenetic marks. We employ a Monte Carlo-based epigenetic labeling exchange algorithm to account for this phenomenon. Since the epigenetically conserved process should maintain a thermodynamic balance and lead to redistribution of only the epigenetic marks, we take inspiration from Kawasaki dynamics, which are used to study spin-conserving Ising systems[66] (Fig. 1g). The steps of our exchange algorithm are as follows:

1. Choose a random chromatin bead i and a neighboring bead j (within 1.8*σ*).
2. The energy change ΔE associated with swapping the epigenetic marks of beads i and j is calculated. The energy change is given by the difference in energy before and after the epigenetic labeling swap.
3. If ΔE ≤ 0, accept the swap, and update the chromatin labeling configuration by exchanging the epigenetic labels at sites i and j.
4. If ΔE > 0, accept the swap with a probability P = exp(−ΔE/kT_E_), where k is the Boltzmann constant and T_E_ is the temperature associated with this exchange. This probability, known as the metropolis criterion, ensures that the system tends to lower its energy over time in a canonical ensemble setting.

To ascertain that such dynamics will not give rise to out-of-equilibrium components and lead the system to a thermodynamically favored state, we show that it follows detailed balance in SI1.4.

#### Epigenetic reactions

To account for changes in the concentration of nuclear epigenetic remodelers, which can be driven by the microenvironment[1, 26, 27, 30], we employ epigenetic reactions. Epigenetic reactions exhibit active dynamics, act in a nonconservative manner and can thus change the net heterochromatin-to-euchromatin (red-to-blue) ratio in the system. Heterochromatin can be converted to euchromatin through the activity of histone demethylases followed by the action of histone acetyltransferases. Here, we quantified the conversion of heterochromatin to euchromatin because of the combined effect of these reactions on the acetylation rate (Γ_*ac*_). Similarly, we quantified the conversion of euchromatin to heterochromatin through the methylation rate (Γ_*me*_). These reactions are modeled as stochastic processes in which a random bead is chosen and converted to heterochromatin or euchromatin bead with rates equal to Γ_*me*_ (methylation) or Γ_*ac*_ (acetylation), respectively (Fig. 1h). When a reaction move is executed, the epigenetic marking of a chosen bead is changed in a probabilistic manner, i.e., the bead changes to blue with probability *p*_*ac*_(Γ_*ac*_) and to red with probability *p*_*me*_ (Γ_*me*_) such that *p*_*ac*_(Γ_*ac*_) + *p*_*me*_ (Γ_*me*_) =1. We show that epigenetic reactions break the detailed balance and introduce out-of-equilibrium dynamics in SI1.4.

The Brownian diffusion timescale (*τ*_*Br*_) is taken as the reference timescale for the system. Additionally, we execute one epigenetic diffusion move with every Brownian step, which gives us an effective diffusivity ∼1*μm*^2^/*s* [67, 68](details in SI1.3). To execute the reactions, we execute one epigenetic reaction step for every 10^3^ simulation steps. Since each simulation step corresponds to 10^−4^s, the reaction rate corresponds to a timescale of ∼0.1 s^−1^, which is in the range of experimentally observed histone remodeler activity [69]. To perform production runs and simulate real-time chromatin organization changes, we first constructed a polymer from experimentally observed sequencing data, as detailed in section 2a. Thereafter, we subject it to diffusion and reaction dynamics.

### 2c Preparation of cells and TSA treatment

A-375 (CRL-1619) cells were procured from ATCC. A-375 cells were isolated from a 54-year-old female patient with malignant melanoma. They possess an epithelial morphology. The cells were resuspended in basal growth medium (DMEM and 10% fetal bovine serum (FBS)). Experiments were performed on cells at greater than 3 passages or fewer than 8 passages. Splitting was performed only after a confluence of more than 80% was achieved.

We used tricostatin A (TSA) as an HDAC inhibitor (Fisher Scientific 14-061). For TSA treatment, the cells were grown to >70% confluence for more than 24 hours before the addition of 500 nM TSA along with basal media. These were returned to the incubator and cultured for another 2 hours. Thereafter, they were washed four times with PBS for 5 min each and then fixed with 4% paraformaldehyde (PFA) in PBS for 10 min (imaging) or 1% formaldehyde in 1x HBSS (genomic assays).

For immunostaining and confocal imaging, after incubation, primary antibody (H3K9ac) diluted in antibody dilution buffer (4% BSA, 0.25% Triton in PBS) was added, and the samples were incubated overnight at 4°C. After primary antibody incubation, the cells were washed 3 times with PBS for 5 minutes each. The cells were then incubated with secondary antibodies (either Alexa Fluor 594 or Alexa Fluor 488) per the manufacturer’s directions for 30 minutes at room temperature. After secondary antibody incubation, the cells were washed 3 times with PBS and sealed with mounting media supplemented with DAPI. Slides were treated with mounting media for 24 hours before imaging. The cells were imaged via a Leica Sp8 confocal microscope equipped with a 63x oil immersion objective.

### 2d STORM imaging and analysis

To perform STORM imaging of the fixed sample, the cells were incubated in blocking buffer containing 10% wt/vol BSA (Sigma) in PBS for 1 h. The samples were then incubated overnight with rabbit anti-H2B (1:100; Proteintech, 15857-1-AP) at 4 °C. Thereafter, the samples were repeatedly washed in PBS, and secondary antibodies (Alexa Fluor 647) were added for imaging. The images were taken on a commercially available ONI (Nanoimager S) STORM microscope system. To ensure optimal photo-switching of Alexa Fluor 647, the imaging buffer followed standard guidelines[70] and consisted of 10 mM cysteamine MEA in Glox solution: 0.5 mg ml-1 1-glucose oxidase, 40 mg ml-1 1-catalase and 10% glucose in PBS. A 640 nm laser was used at a setting of 40% to excite the reported dye (Alex Fluor 647), and a gradual increase in the 405 nm laser was used to reactivate Alexa Fluor 647 in an activator dye to maintain a constant intensity of active fluorophores. An exposure time of 15 ms and 30k frames was used. Downstream analysis was performed with the help of a custom code in MATLAB (SI3.1), which we showed in a previous publication [14].

### 2e Hi-C preparation

Biological replicates of Hi-C were performed on two independent samples of A375 cells, both untreated and treated with 0.5 µM TSA for 2 hr. The first biological replicate was processed with the protocol described previously[71]. Briefly, 10 million cells were crosslinked with 1% formaldehyde for 10 min. Crosslinked cells were then suspended in lysis buffer to permeabilize the cell membrane and homogenized. Chromatin was then digested in the nucleus overnight using the DpnII restriction enzyme. The digested ends were filled with biotin-dATP, and the blunt ends of the interacting fragments were ligated together. The DNA was then purified via phenol‒ chloroform extraction. For library preparation, the NEBNext Ultra II DNA Library prep kit (NEB) was used for libraries with sizes ranging from 200–400 bp. End Prep, Adaptor Ligation, and PCR amplification reactions were carried out on bead-bound DNA libraries. The second biological replicate was processed via the Arima-HiC+ Kit from Arima Genomics following the protocol for Mammalian Cell Lines (A160134 v01), and the libraries were prepared via the Arima recommendations for the NEBNext Ultra II DNA library prep kit (protocol version A16041v01).

Sequencing was performed on a HiSeq 3000 platform with 150 bp paired-end reads. Sequencing reads were mapped to the human genome (hg19), filtered, binned, and iteratively corrected as previously described (Imakaev et al., 2012) via the HiCPro pipeline (https://github.com/nservant/HiC-Pro). Spatial compartmentalization (A/B compartment assignment) was calculated via principal component analysis with 40 kb binned data.

### 2f RNA-seq preparation

RNA was extracted from TSA-treated and untreated A375 cells via the Qiagen RNeasy Plus Mini Kit (Cat. No. 74134). The cells were lysed, homogenized, and spun down in gDNA eliminator columns to remove any genomic DNA. All the samples were washed with ethanol, and the total RNA was eluted. The quality and quantity of the RNA were quantified via an Agilent Bioanalyzer Nano RNA Kit. All the libraries used were characterized by an RNA integrity number (RIN) between 9 and 10. The RNA was sent to the Oklahoma Medical Research Foundation for library preparation (using an rRNA depletion approach) and sequencing.

The quality of the reads was checked via FastQC, and adapter sequences were trimmed via BBTools (https://github.com/kbaseapps/BBTools). Additionally, quality trimming of the reads was performed, and any reads with a quality score lower than 28 were discarded. The reads were then aligned via STAR alignment (https://github.com/alexdobin/STAR). The aligned reads were then sorted by genomic position, and feature counts were performed via HTSeq 0.11.1 (https://github.com/simon-anders/htseq). The differential expression of genes was determined via DESeq2[72].

## 3 Results

### 3a Interplay of diffusion and reaction dynamics drives chromatin domain formation and size scaling

We start by quantifying the 3D spatial organization principles, defining the nanoscale behavior of the chromatin polymer model through analysis of its temporal evolution. We illustrate how diffusion and reaction dynamics (as described in section 2b) work in tandem to create nanoscale heterochromatic domains, resulting in a scaling relationship between domain size and the rates of diffusion and epigenetic reactions. To distinguish this physical phenomenon from those that depend on specific details of heterochromatin and euchromatin segment arrangements, we first investigate the prototype random polymer model (described in section 2a and SI1.1). For the prototype polymer, we choose the heterochromatin-to-euchromatin ratio to be 1.5, mirroring the prevalent observation in most Hi-C maps (SI1.1).

#### Diffusion of epigenetic marks leads to Ostwald ripening

We start by considering only spatial (Brownian) diffusion of chromatin and the diffusion of epigenetic marks. This ensures that the system adheres to thermodynamic balance, and the global ratio of active (blue) and repressed (red) chromatin segments (beads) remains constant throughout the simulation. Commencing with the prototype polymer in a random configuration, as illustrated in the left panel of Fig. 2a, we track the temporal evolution by monitoring the radius of gyration of the heterochromatin-rich domains. The methodology for calculating the radius of gyration is described in SI1.5. As the system evolves, it reaches a steady-state configuration (Fig. 2a, red simulation trajectory), which remains constant over time. Upon closer examination of the trajectory, we observe initial clustering and enlargement of the repressed regions facilitated by the diffusion of epigenetic marks, as quantified by the initial growth in the radius of gyration of the repressed domains. A complete spatial segregation of the epigenetic marks subsequently occurs, with the repressed marks migrating toward the center of the simulation box while active marks are displaced outward, depicted through the plateauing of the radius of gyration. This spatial segregation represents a thermodynamically favored state of the system, which minimizes the interfacial area between the active and repressed phases, reminiscent of Ostwald ripening. These simulations demonstrate that passive diffusion alone fails to prevent the coarsening of epigenetic marks and thus does not lead to the formation of characteristic chromatin domains.

**Figure 2:**
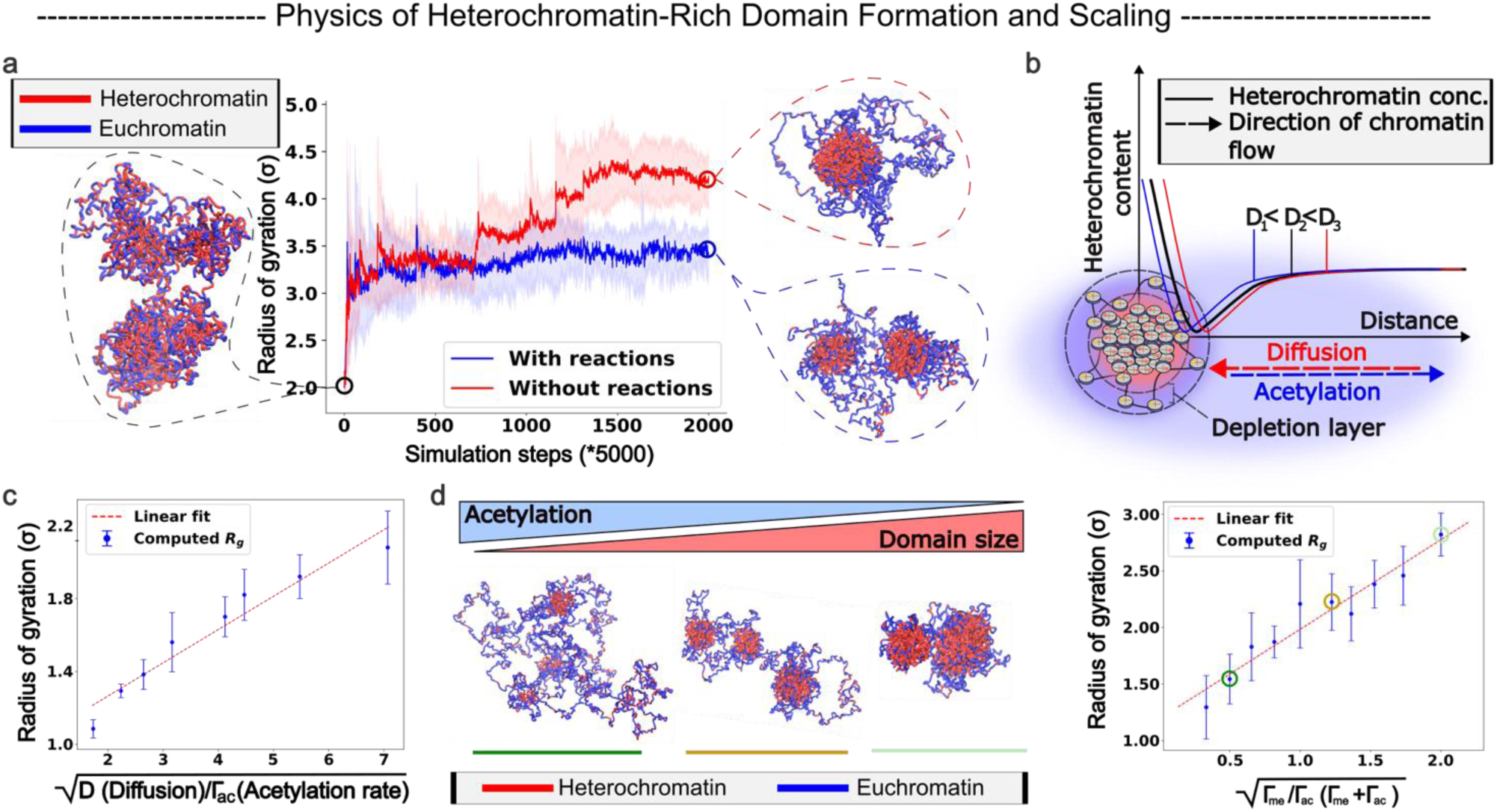
Chromatin dynamics form finite-sized domains and exhibit domain scaling. a) The evolution of the radius of gyration of the domains of the prototype polymer in the presence and absence of epigenetic reactions. Starting from a randomly labeled (60% heterochromatin) polymer, the proposed dynamics lead to ripening of the heterochromatin domain in the absence of reactions and readily form finite-sized domains in their presence. b) Schematic showing the action of diffusion and acetylation on the heterochromatin packing domains. The y-axis shows the heterochromatin concentration, and the x-axis shows the distance from the center of the domain. Diffusion and acetylation act in opposite directions, with diffusion leading to domain growth and acetylation to shrinkage. The empirical heterochromatin content vs. distance plots have been provided for three different diffusion rates (D) with the same reaction rates. c) A scaling relationship is observed between the radius of gyration and the relative rates of diffusion and reactions, which dictates that the radii of the domains scale linearly with 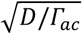. d) With constant diffusion, as the reaction rates change, the domain sizes change. Higher methylation (lower acetylation) drives an increase in domain size (polymer snapshots). A scaling relation is obtained, which shows that the average radius of gyration of the heterochromatin domains is directly proportional to 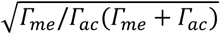 (right). Circled points on the plot represent the parameters for the polymer configurations on the left.

#### Epigenetic reactions compete with diffusion to give rise to domains of a characteristic size

Next, we introduce epigenetic reactions into our system. These reactions lead to out-of-equilibrium interconversion of epigenetic mark dynamics, which can alter the global ratio of repressed (red) and active (blue) polymer beads (details on detailed balance considerations in SI1.4). Notably, in the previous scenario where no reactions were present, the net heterochromatin-to-euchromatin ratio remained the same as that of the initial polymer. However, with the introduction of reactions, this ratio becomes variable and stabilizes at a steady state equal to the ratio of the rates of the methylation and acetylation reactions. In this context, we maintain the reaction ratio identical to the initial polymer composition to facilitate an unbiased comparison with the diffusion-only scenario, i.e., Γ_*me*_ /Γ_*ac*_ = 1.5. The initial polymer configuration is also the same as that of the diffusion-only setting (Fig. 2a). Like in the diffusion-only case, we observe an initial ripening phase where the repressed marks diffuse to form phase-separated domains (Fig. 2a, blue simulation trajectory). However, as the simulation progresses, the growth of these domains levels off, resulting in distinct and persistent separated domains. These repressed domains exhibit a characteristic length scale akin to what is observed in super resolution imaging studies[1, 23, 24]. Moreover, the simulation illustrates that these repressed domains maintain their characteristic shapes over an extended period (∼10 hours of real-time simulation), comparable to the timescale of a typical duration of human cell interphase. This suggests that by incorporating the out-of-equilibrium dynamics of epigenetic reactions alongside equilibrium-preserving diffusion dynamics, the simulation yields heterochromatin domains that are stable against Ostwald ripening.

To understand the biophysical underpinnings of domain formation in more detail, we next elucidate the underlying mechanisms. During the initial growth phase of the domains (Fig. 2a), favorable interactions facilitate the clustering of similar epigenetic marks through energetically driven Brownian diffusion and the diffusion of epigenetic marks. This aggregation leads to the formation of heterochromatin-rich domains surrounded by heterochromatin-poor regions (Fig. 2b). The depletion of heterochromatin from the domain’s immediate vicinity establishes a flux of heterochromatin regions toward the formed domain, driving its growth. In the absence of reactions, this flux leads to complete phase separation. However, when reactions are introduced, the acetylation reaction (as quantified by Γ_*ac*_), which converts heterochromatin to euchromatin in a spatially invariant manner, converts heterochromatin (red) to euchromatin (blue) within the growing domains, competing with the diffusion process. As the simulation progresses, the reaction kinetics hinder complete phase separation. Domain growth ceases when the influx of heterochromatin through diffusion is balanced by the conversion of heterochromatin to euchromatin within the domain via the acetylation reaction. A scaling argument (SI1.6) shows that the radii of the heterochromatic domains *R*_*d*_ follow the scaling relation [1]:

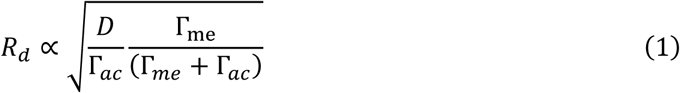

where *D* is the diffusion rate of epigenetic marks and Γ_*me*_ and Γ_*ac*_ are the methylation and acetylation rates, respectively (as explained in section 2b). To ascertain this diffusion‒reaction relationship through our proposed MD model, we fix the reaction rates and vary only the diffusion rates in our simulation (Fig. 2c). The simulation confirms the nonlinear relationship of the radius, which increases with diffusion and is counteracted by acetylation, i.e., proportional to 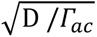. This finding establishes a central biophysical concept through our polymer model - reactions and diffusion compete to give rise to chromatin domains - a result that has not been previously reported in a molecular dynamics setting.

#### Chromatin domains exhibit nonlinear scaling with changing epigenetic reaction rates

Recent advancements have highlighted the dynamic regulation of chromatin organization in response to chemo-mechanical cues from the cellular microenvironment, which influences the nuclear concentration of key epigenetic modifiers, mainly the histone writers and erasers [14–16]. The resulting epigenetic reactions, which capture shifts in the nuclear concentration of these remodelers, dictate the overall chromatin landscape by altering the heterochromatin-to-euchromatin ratio within the nucleus. The global effect of these reactions on chromatin organization is captured in eq (1), which we validate here. For this analysis, we fix the diffusion constant, focusing on understanding how the balance between methylation and acetylation reactions drives chromatin reorganization. We quantified the radius of gyration of repressed chromatin domains as a function of epigenetic reaction rates to gauge the extent of chromatin reorganization (Fig 2d). As illustrated in Fig. 2d, the radius of gyration decreases as the acetylation rates increase (or the methylation rates decrease), confirming the analytical relationship in eq (1). Thus, we establish that as the number of acetylation (methylation) marks increases, there is a greater tendency for chromatin decompaction (compaction). This alteration in compactness can consequently affect chromatin accessibility in specific regions. The modified epigenetic marking and accessibility of chromatin alters potential interactions with the transcriptional machinery, thereby influencing gene expression. Hence, our model links changes in epigenetic modifier concentrations to chromatin organization and downstream effects via a single dimensionless factor—the ratio of the methylation rate to the acetylation rate (Γ_*me*_ /Γ_*ac*_)—and demonstrates how the sizes of chromatin domains change with this factor.

### 3b Data-informed chromatin polymer predicts spatial and epigenomic alterations in response to changing epigenetic reactions

We initialize our simulations with experimentally derived epigenetic data, a critical step that significantly influences the subsequent chromatin reorganization dynamics. This represents a classic initial value problem, where the initial epigenetic state dictates the cell’s epigenetic trajectory, shaping its future chromatin conformation and gene expression profile. Our model’s robust generalization, rooted in its biophysical foundations as demonstrated in the preceding section where domains emerge in the prototype polymer, enables it to leverage sequencing sources such as ChIP-seq or Hi-C to guide the initial epigenetic distribution. As described in section 2a, we utilize Hi-C data for A375 cells (malignant melanoma with epithelial morphology) and ChIP-seq data for human mesenchymal stem cells (hMSCs derived from human bone marrow) in the following sections. Our objective is to bridge the observed chromatin reorganization across mesoscale imaging and genomic sequencing, identifying potential genomic loci that are altered in response to changes in epigenetic reactions or the microenvironment.

In this section, by simulating the A375-informed chromatin polymer, we elucidate two key phenomena via our model: (i) the stability of chromatin epigenetic distribution in control conditions and (ii) the identification of genomic loci where epigenetic changes predominantly occur in response to changes in these rates.

#### The data-informed polymer model preserves the chromatin domains with constant reaction rates

To ensure that changes in chromatin organization are preserved under steady-state conditions, we examine the stability of chromatin domains when the reaction rates remain constant. Here, stability encompasses the epigenetic labeling of genomic loci and the spatial distance between chromatin segments, collectively influencing the cell’s gene expression profile. The reaction ratio, Γ_*me*_/Γ_*ac*_, is chosen to be the same as the initial heterochromatin-to-euchromatin ratio. The temporal kymograph in Fig. 3a (chromosome 19 of A375) shows the epigenetic marks at each chromatin segment (bead) as the simulation progresses. Remarkably, the initial distribution of epigenetic marks largely persists throughout the simulation, retaining its identity over a span of 2 hours of real time (extended simulations up to 10 hours are illustrated in SI2.1), which demonstrates the ability of our model to maintain the epigenetic identity of individual chromatin segments. Additionally, we present the segment-to-segment distance matrix of the polymer at the start and end of the same simulation in SI2.1. The alignment between the initial and final pairwise distances of the chromatin segments further confirms the stability of the chromatin configuration as influenced by its dynamics. Thus, our model not only preserves the epigenetic identity but also maintains the accessibility of chromatin segments. Analogous analyses have been conducted for chromosomes 18, 20, and 21 in SI2.1.

**Figure 3:**
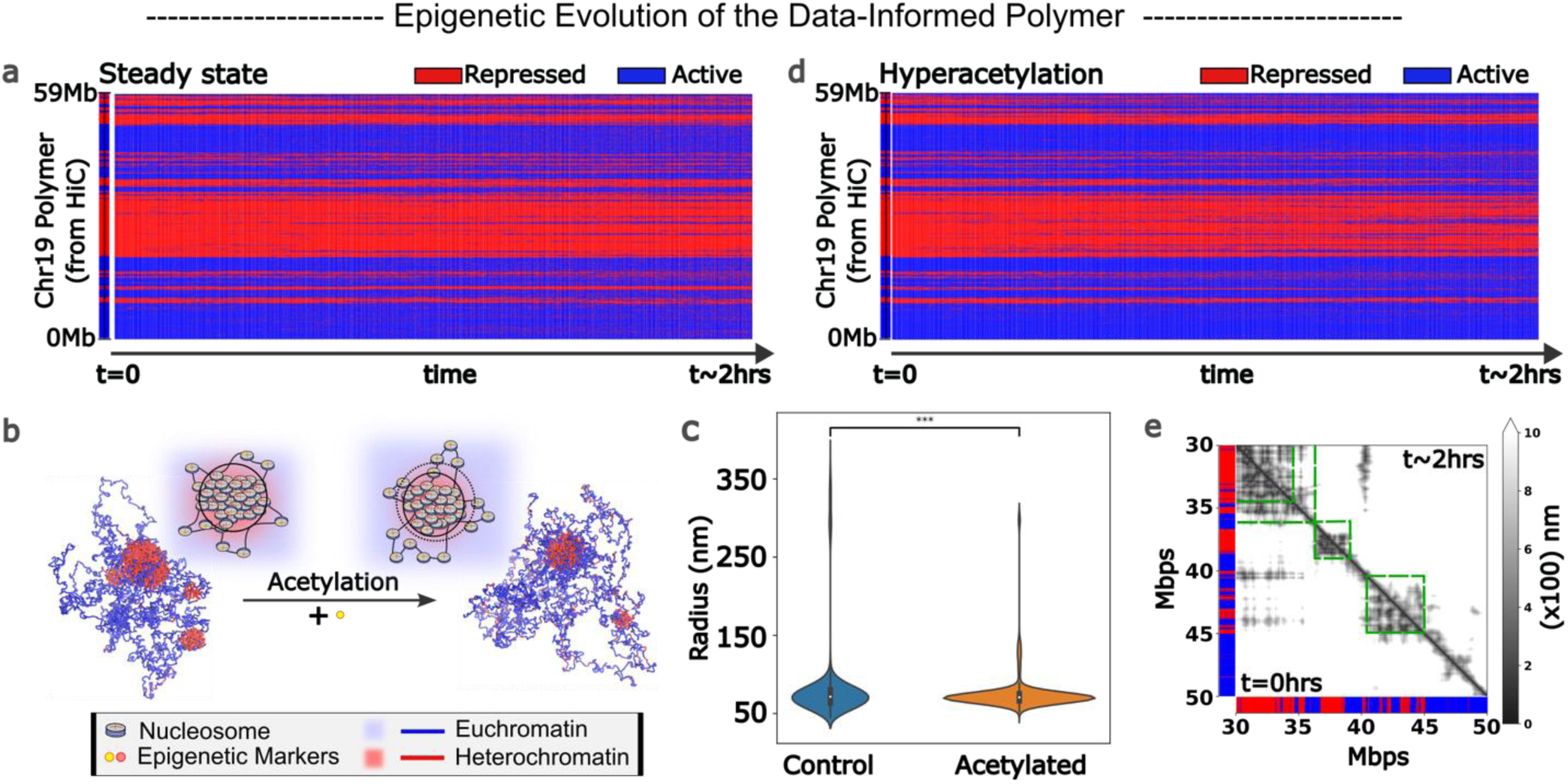
Effects of changing the concentrations of histone epigenetic remodelers on chromatin. a) Temporal kymograph of chromosome 19 of A375 cells when the ratio of methylation to acetylation is the same as the initial heterochromatin to euchromatin ratio. The epigenetic domains temporally maintain their genomic extent on the polymer. b) With increasing acetylation, the repressed domain sizes decrease, as observed through the simulation snapshots. This finding was further validated through the observed radius of gyration of the chromatin domains in c). d) Kymograph evolution exhibiting euchromatin domains spread out to neighboring genomic regions as the number of acetylated beads increased. e) The pairwise distance map shows the spatial reconfiguration of the polymer upon acetylation. The lower triangle corresponds to the initial pairwise distance between the polymer beads, and the upper triangle corresponds to the final pairwise distance. Upon acetylation, major decompaction is observed at the boundaries of heterochromatin domains, as shown by the increasing pairwise distance in the green boxes.

#### Modifying the reaction rates alters epigenetic marks at the boundaries of domains

We next examine the changes in epigenetic marks of the polymer segments in response to variations in the reaction rates. We simulate a scenario where the acetylation reaction rate increases, resulting in a lower methylation-to-acetylation ratio (0.2) than the initial heterochromatin-to-euchromatin ratio (∼0.45) for 2 hours in real time. This alteration in the reaction ratio led to a decrease in the size of the repressed chromatin domains, as evident from the reduced radius of gyration of these domains in Fig. 3b, which was further quantitatively confirmed in Fig. 3c. However, specific loci along the 1D polymer that exhibit alterations in epigenetic marks and accessibility as chromatin undergoes a transition to a more acetylated state can be identified in the case of data-informed polymers. The temporal kymograph of the polymer evolution (Fig 3d) shows that the switch from the repressed to the active state is prominently localized to the domain boundaries of larger chromatin domains (∼100s of kbps) or smaller chromatin domains (few kbps) on the polymer. Our prediction shows that this heterochromatin-to-euchromatin conversion at the boundaries leads to chromatin decompaction, which is evident in the pairwise distance map (Fig. 3e). Since boundary regions undergo epigenetic and spatial changes, these changes can potentially affect the transcriptional activity of genes at these genomic loci. We extended this analysis to other chromosomes in the A375 cell line, as presented in SI2.2. In SI2.2, we also show that an increased methylation rate changes genomic loci from euchromatin to heterochromatin, leading to chromatin compaction, which is also predominantly observed at domain boundaries.

This phenomenon of epigenetic marking alterations predominantly at domain boundaries arises from the fact that heterochromatin–heterochromatin interactions are significantly stronger than heterochromatin–euchromatin interactions, making it energetically favorable for epigenetic changes to occur at boundaries rather than within heterochromatin-rich domains, where they would require breaking favorable interactions. In conclusion, our dynamic model reveals a fundamental principle: chromatin regions at domain boundaries are most susceptible to changes in the concentration of epigenetic remodelers.

### 3c Polymer model integrates complementary observations from super-resolution imaging and Hi-C sequencing, revealing epigenetic change specificity at domain boundaries

Next, we utilized super-resolution imaging (STORM) alongside bulk or population-level Hi-C sequencing to probe chromatin organization. Integrating these methods presents a challenge, as they offer complementary yet partial insights: STORM reveals the nanoscale organization of chromatin domains but lacks specific genomic loci data, whereas Hi-C supplies contact details for individual genomic loci but with limited spatial information. Furthermore, bulk techniques such as Hi-C overlook cellular heterogeneity, a factor captured by imaging. Our data-informed polymer model can predict both 3D morphological changes and associated genomic loci, thereby bridging this gap, and offering a comprehensive understanding of chromatin organization. We conducted super resolution STORM imaging and bulk Hi-C sequencing on melanoma cells subjected to identical epigenetic modifications and analyzed the results with our polymer model to link these complimentary methods. This approach enabled us to elucidate the concomitant changes occurring at both the mesoscale (∼100s of kb scale) and the genomic scale (kb scale), providing a multiscale understanding of chromatin reorganization in response to changes in epigenetic reaction rates.

To investigate the effects of changes in epigenetic reaction rates on chromatin organization, we treated A375 melanoma cells with 0.5 μM trichostatin A (TSA), a well-characterized HDAC inhibitor, for 2 hours. This treatment is expected to increase acetylation levels, leading to a more decompacted, euchromatin-rich chromatin organization. We chose a 2-hour timepoint to allow sufficient time for histone modifications to occur without secondary effects such as transcriptional changes influencing chromatin organization[73]. Immunofluorescence imaging of H3K9ac revealed a global increase in acetylation levels (Fig. 4a), while DAPI staining revealed significant changes in chromatin reorganization (Fig. 4a). To examine these changes at higher resolution, we performed STORM imaging, as described in the following subsection.

**Figure 4:**
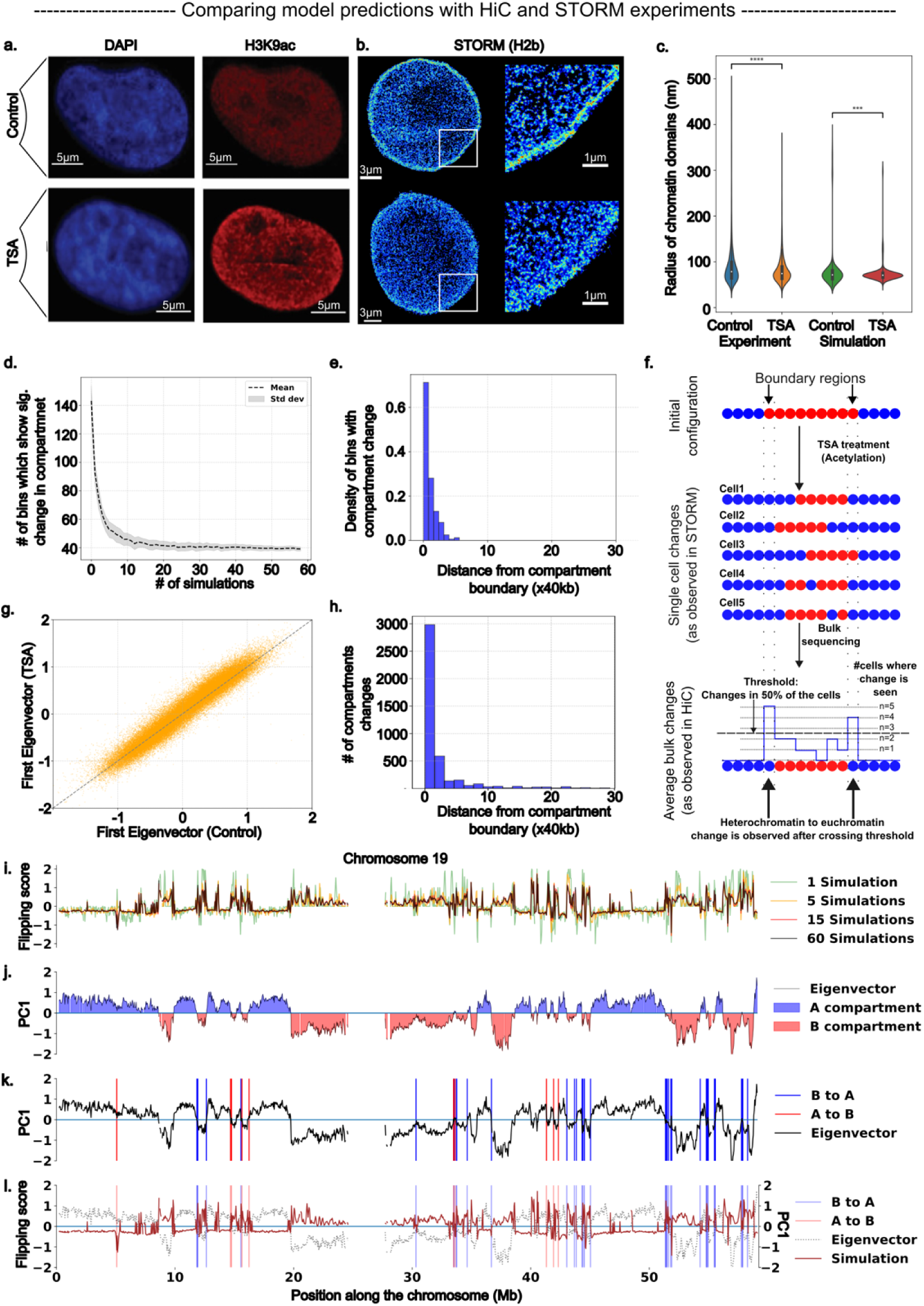
Chromatin polymer model bridges STORM and Hi-C observations: a) TSA treatment of A375 melanoma cells shows chromatin reorganization (DAPI, blue) and increase in nuclear acetylation (H3K9ac, red). b) H2b labeled STORM images display the contrast in chromatin organization between control and treated cells. TSA treated cells show chromatin decompaction with smaller interior chromatin domains. c) Simulation captures trends of the changes observed in chromatin domain radii through STORM. A higher acetylation rate is used to replicate the hyperacetylated A375 nuclei. d) To match our simulations with bulk sequencing, we average our compartmental changes over multiple simulations to capture the effect of combining multiple single cell epigenomics to get the population average. For chr19 of A375 cells, the number of 10 kb beads which show a significant compartment change falls by threefold after we average 30+ simulations and plateaus after that. (e) Most of the chromatin segments (beads) which change their epigenetic marking in simulations lie very close to the domain boundaries. (f) The schematic shows the differences observed in single cell vs bulk sequenced samples. Each individual cell may have a heterochromatin domain decrease from 9 to 5 beads, but, in the bulk averaged setting, the domain size we obtain is 7 beads. This shows that bulk averaging exhibits the changes observed in majority of the cells and masks cell-to-cell heterogeneity. g) The PC1 of Hi-C(TSA) vs Hi-C(Control) (40 kb bins) shows little change in the treated Hi-C compared with the control. h) The frequency of bins with changed compartments vs. distance from the closest boundary shows that the compartmental changes observed in the Hi-C across the genome are very close to the domain boundaries, with 70% within 40 kb of the closest domain boundary. i) The simulation predictions on compartment changes for chr19 of A375 are plotted as a flipping score along the genome. This flipping score is normalized to 2, with 2 (−2) showing a perfect probability to change compartment from B to A (or A to B for negative values). The absolute flipping scores decline and then stabilize as simulations. We deem a flipping score greater than 1 or −1 to constitute a significant flip. j) A and B compartment profile as obtained from the PCA of control Hi-C contacts for chr19 of A375 cells. k) Compartment changes as observed from the control and Hi-C maps. B to A (blue vertical lines) mark the bins where a compartment change from B to A is observed and similarly for A to B is shown with red lines. l) Overplotting i) and k) shows good agreement of the predicted and observed compartment changes.

#### STORM imaging confirmed chromatin domain size scaling in agreement with model predictions

Super-resolution STORM imaging confirmed the presence of characteristic chromatin domains in A375 melanoma cells (Fig. 4b). Notably, following TSA treatment, the domain sizes significantly decreased across the two replicates (∼5 cells each), as depicted by the representative STORM images in Fig. 4b. We quantified the domain sizes (detailed in SI3.1) across replicates and observed a statistically significant decrease in domain size (Fig. 4c). To validate these changes with our model predictions, we conducted simulations over a 2-hour real-time span, replicating the TSA treatment conditions by increasing the acetylation rate. These simulations were performed on chromosomes 18-21 of A375 cells. The simulated control and acetylated chromatin domains, as predicted by the simulation, demonstrated a marked decrease in the observed radius of gyration of the chromatin domains. The mean of the experimentally observed domain radii decreases by 8%, aligning closely with the simulation prediction of an 11% reduction. Furthermore, the size distributions of the experimental and simulated domains before and after operation are similar, as illustrated in Fig. 4c.

#### Averaging over the cell population leads to fewer epigenetic compartmental changes

To explore the relationship between single-cell STORM and bulk Hi-C sequencing, we integrated population and single-cell effects into our simulations. At the single-cell level, kymographs illustrating the simulated effects of chromosome 19 acetylation in A375 cells (Fig 3d) demonstrated domain shrinkage, suggesting a preference for compartmental shifts toward a more euchromatic configuration at domain boundaries. These findings indicate that epigenetic boundary regions are primarily responsive to changes in epigenetic remodeler activity. To quantify population-level changes, we performed multiple simulations with diverse initial spatial configurations of the chromatin polymer while maintaining Hi-C-informed epigenetic distributions. We introduced a flipping score to measure alterations in epigenetic marks, establishing a threshold to identify regions with significant changes (further elaborated in SI4.1). The flipping score assesses the proportion of cells exhibiting a compartmental shift, with the threshold set to detect significant changes present in the majority (>50%) of observed cells. This approach facilitates the integration of cellular heterogeneity into our analysis, enabling us to consider its influence on population-level observations. As a result, we gain a more nuanced understanding of chromatin organization and its variability across cells.

As we averaged over simulations, fewer regions passed the threshold for compartment change (Fig. 4d, 4i), demonstrating how cellular heterogeneity is reflected in bulk experiments. As presented in Fig. 4d and 4i, averaging over multiple simulations leads to the suppression of domain changes, which are specific to only a subpopulation (corresponding to averaging over a few simulations), and reveals only those changes that are present more frequently in the whole population (which corresponds to averaging over multiple simulations). Notably, chromatin segments with high flipping scores at domain boundaries survived such an averaging operation (Fig. 4e), as they corresponded to changes in most of the cells. We elucidate this general observation through the schematic in Fig. 4f, which illustrates that heterogeneous cell-to-cell differences are more prominent in single-cell observations than in bulk sequencing observations, where population averaging obscures these variations. The schematic highlights how population-wide changes, such as those at the boundary domains that are present for most of the cells, are captured in the Hi-C observations (or other bulk sequencing observations), but other subpopulation-level changes are not. Our analysis highlights the importance of considering cellular heterogeneity in single-cell vs. bulk experiments, as bulk experiments such as Hi-C sequencing good at highlighting the significant population-wide changes. To validate our genomic-scale predictions, we turn to bulk Hi-C experiments, which provide a means to robustly quantify compartment changes.

#### Changes in bulk Hi-C contact maps reveal epigenetic compartment shifts at domain boundaries

We generated Hi-C maps for two replicates of control and TSA-treated A375 cells, revealing only minor rearrangements (SI4.1) despite substantial alterations in domain size scaling under STORM imaging and disparate H3K9ac immunofluorescence levels. To systematically assess changes in the Hi-C maps, we combined the two Hi-C matrices at 40 kb resolution and employed principal component analysis (PCA) to delineate A (active, euchromatin-like) and B (inactive, heterochromatin-like) compartments for both control and TSA-treated cells (Fig. 4j) (details on Hi-C analysis in SI4.1). Comparative PC1 analysis revealed that most regions retained their compartment identity, with no significant shifts in epigenetic patterns (Fig. 4g). Among approximately 70k mappable bins at 40 kb resolution, only about 5k (∼7%) bins exhibited changes in genomic compartments. Notably, among these 5k bins, nearly 3.1k bins switched from B to A compartments, whereas ∼1.3k shifted from A to B compartments. We defined bins that transitioned from negative to positive PCA values as those that shifted from B-like to A-like behavior, and vice versa. This trend persisted when a threshold based on compartment strength was applied (TSA PC1 < −0.01 for bins going from A to B and TSA PC1 > 0.01 when going from B to A), yielding 2838 bins switching from B to A and 1148 bins switching from A to B. Although these changes are relatively small in the context of the entire genome, they affirm our key predictions: regions tend to transition from repressed to active states upon TSA treatment. The observation that more noticeable changes in STORM imaging correspond to smaller changes in Hi-C contact patterns confirms our previous discussion on bulk and single-cell observations. Our model clearly shows that significant changes in single-cell STORM imaging correspond to subdued changes in Hi-C data due to averaging. Next, we analyzed the precise genomic locations of these compartment switches.

We examined the initial epigenetic characteristics of the compartments that underwent state changes, revealing that nearly half of the bins that shifted compartmental signatures had initial PC1 absolute values of 0.1 or less. These findings suggest that these regions are initially less strongly associated with their specific epigenomic domains, increasing their susceptibility to alteration upon epigenetic perturbation. To investigate the genomic locations of these altered bins, we plotted their positions on the genome. Fig. 4k shows the genomic positions of bins transitioning their epigenetic compartment from A to B (marked by blue vertical lines) and from B to A (marked by red vertical lines) for chromosome 19 of A375 cells. This visual representation reveals that the observed compartmental shifts predominantly occur very close to the domain boundaries. To quantify this result, we analyzed this phenomenon genome-wide, which revealed that most compartment switches (over 75%) are located within 50 kb of boundary regions (Fig. 4h). This finding confirms that compartmental boundaries are notably responsive to changes in epigenetic reactions. When we overlay these experimental changes with the predictions generated by our polymer model after averaging over multiple simulations (Fig. 4l), a clear alignment emerges, with high flipping scores aligning with positions where a compartment change is observed in Hi-C.

In summary, our model highlights two key results: 1) While we observe noticeable changes in STORM imaging, the Hi-C data reflect only those changes that are present for the majority of the cells while not capturing changes that are shown by smaller subpopulations. 2) The compartment boundaries, as confirmed by Hi-C observations, emerge as regions of heightened sensitivity to epigenetic reaction alterations, where transitions in chromatin compartmentalization are favored owing to the enacting kinetics. We next analyze the downstream effects of these epigenetic changes via RNA-seq analysis.

### 3d. RNA-Seq of A375 cells revealed key differentially expressed genes near domain boundaries that are pivotal for epithelial-to-mesenchymal transition and metastasis

To analyze the changes in gene expression that accompany chromatin reorganization, we performed RNA-seq on 2-hour TSA-treated A375 cells. Among the significantly differentially expressed (DE) genes (|*l*og2(TSA/Control)| > 1 and pvalue < 0.05), 423 genes were upregulated, and 124 genes were downregulated (Fig 5a). This finding is consistent with our model prediction that HDAC inhibition leads to increased euchromatin content and thus can cause preferential global upregulation of genes. Analysis of the RNA-seq data with clusterProfiler[74] via gene ontology (GO) overrepresentation analysis (ORA) for biological pathways revealed that among the differentially regulated pathways, the most significant were those related to cell migration (Fig 5b, top panel). Since A375 cells are malignant melanomas with an epithelial morphology, this observation suggests an altered metastatic potential for cells treated with TSA[75]. We also found that the Wnt signaling and MAPK/ERK pathways were significantly differentially regulated (Fig. 5b, bottom panel). A large body of published work shows that Wnt (canonical and noncanonical) signaling[76–78] and MAPK/ERK signaling[79–81] are instrumental in driving the epithelial-to-mesenchymal transition (EMT). This could translate to an altered metastatic tendency in A375 cells through EMT[82–88] upon TSA treatment and therefore be differentially regulated with changes in the tumor microenvironment. Our findings further confirm the widely recognized phenomenon that HDAC inhibition leads to altered metastatic potential by differentially regulating key pathways[89–94].

**Figure 5:**
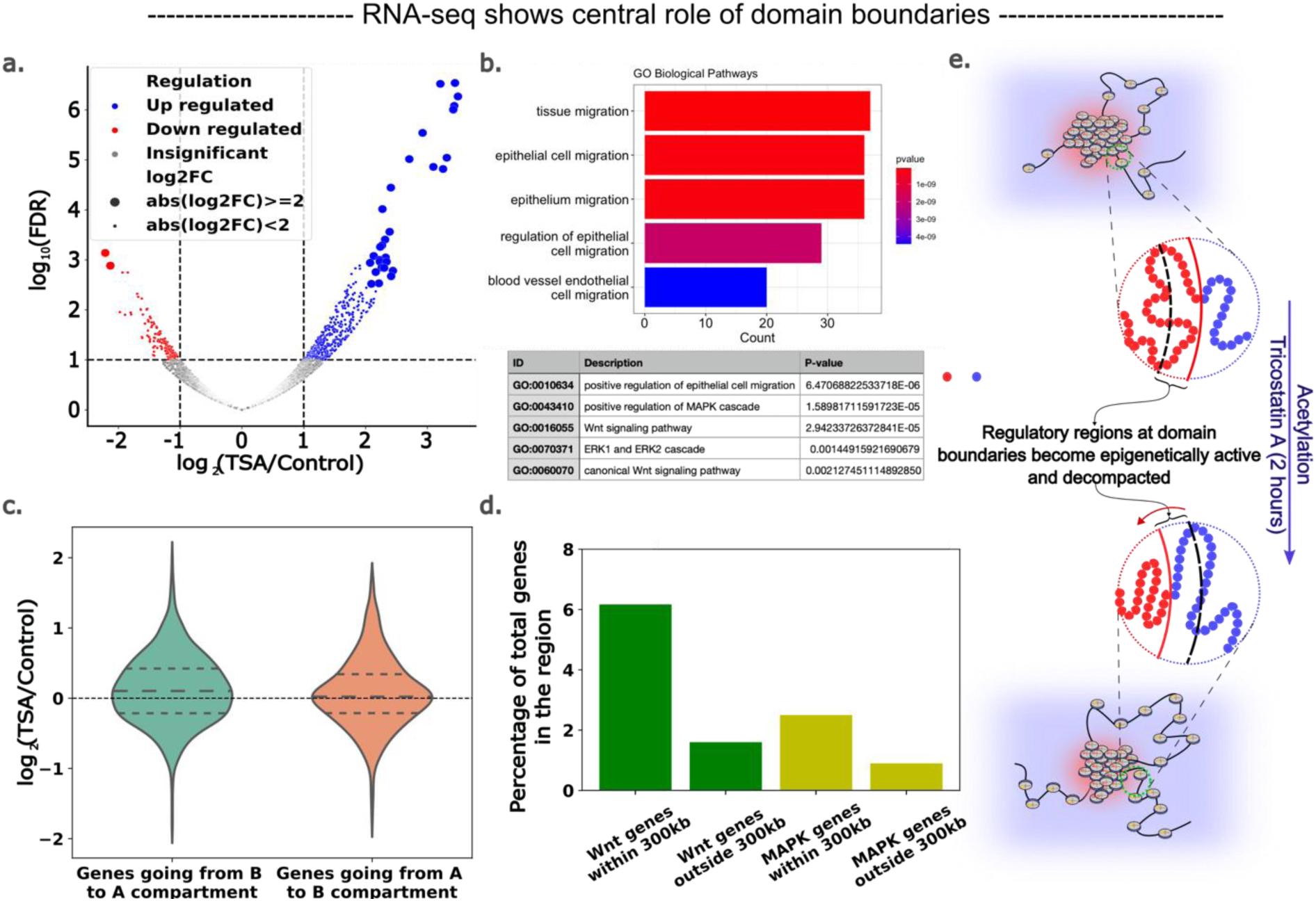
Epigenetic changes at domain boundaries can play a central role in determining cell fate. a) Global RNA-seq trends showing that among the significantly differentially expressed genes, ∼420 genes are upregulated and ∼120 are downregulated, indicating that, upon HDAC inhibition (TSA treatment), genes are preferentially upregulated globally. b) Overrepresentation analysis of biological pathways of the upregulated genes via clusterProfiler shows that cell migration is significantly altered after treatment. Moreover, the canonical Wnt signaling and MAPK/ERK pathways were significantly upregulated. These pathways are particularly important for epithelial-to-mesenchymal transition and are therefore central to metastasis. c) Among all genes belonging to genomic bins where an epigenetic shift is observed in the Hi-C data, genes belonging to the bins with B going to the A compartment show a significant upregulation trend, whereas the opposite compartment flip has no discernable trend. d) Focusing on genes that are differentially regulated from the Wnt and MAPK pathways, we show that they are significantly enriched closer to the domain boundaries than away from it, suggesting that epigenetic changes at the domain boundaries may drive these changes. e) Schematic showing how decompaction and epigenetic transformation at domain boundaries expose regulatory regions. This allows for differential expression of the pathways associated with genes/genes with regulatory elements (enhancer, promoter, etc.) in this region.

We integrated RNA-seq data, model predictions, and Hi-C data by analyzing genes in genomic bins with compartment changes. We segregated all genes into two groups: those in bins transitioning from A to B compartments and those in bins transitioning from B to A (Fig. 5c). Our analysis revealed a significant preference for upregulation among genes transitioning from B to A but no discernible trends for genes transitioning from A to B. This finding supports our model’s predictions, suggesting that hyperacetylation-induced decompaction at domain boundaries enhances gene upregulation. Notably, we found that differentially regulated genes at domain boundaries are highly enriched for positive regulation of epithelial cell migration (GO:0010634, FDR = 3.3e-05), indicating that epigenetic remodeling at domain boundaries may play a key role in regulating this critical melanoma metastasis process.

Since gene regulation can be driven by epigenetic changes several kilobases away from the coding region, we categorized all the differentially expressed genes within 300 kb as boundary genes (∼30% of DE genes) and the remaining genes as genes located away from the domain boundary. We performed GO-ORA analysis for the genes within 300 kb, which revealed that pathways associated with epithelial-to-mesenchymal transition were enriched in this gene set (SI4.2). To ascertain that this is not a general trend among any subset of DE genes and is specific to genes close to the domain boundaries, we compared the enrichment of the EMT gene sets close to the domain boundary against those away from the boundary. Specifically, we focused on EMT-relevant GO pathway subsets (Wnt and MAPK/ERK) that were enriched in the global subset. For these pathways, we plotted their gene ratios within the 300 kb region and outside the 300 kb region (Fig. 5d). We found that both pathways are significantly overrepresented closer to the domain boundary than away from it, suggesting that perturbations to the epigenetic domain boundaries can drive changes in their expression (gene list provided in SI4.2). This analysis revealed that perturbations in nuclear epigenetic remodelers, such as those of HDACs, not only epigenetically drive changes enriched at domain boundaries but also potentially lead to differential regulation of critical genes in these regions.

In summary, this section elucidates the pivotal role of genes near domain boundaries in orchestrating cellular behavior by transitioning between active and inactive states (depicted in Fig. 5e). Our analysis reveals a robust, overarching mechanism for modulating the cellular phenotype through chromatin reorganization. Importantly, this mechanism facilitates cell line- or history-dependent alterations in cellular behavior resulting from the significant influence of preexisting epigenetic landscapes on phenotypic state changes. Thus, in addition to specific transcriptional pathways governing gene networks, this global mechanism can engender cellular heterogeneity based on a cell’s developmental history. While our demonstration focuses primarily on these changes in response to TSA treatment, we broaden the scope to underscore the wider applicability of our model and demonstrate the versatility of the framework we have established.

### 3e Polymer model predicts chromatin domain size distribution after microenvironmental stiffness changes

In the previous sections, we established the occurrence and scaling of the chromatin domains as predicted by the polymer model in response to changing nuclear epigenetic remodeler concentrations and established our findings through HDAC inhibition via STORM and Hi-C sequencing. However, nuclear chromatin remodeler concentrations can also change in response to changes in the mechanical properties of the microenvironment. Specifically, it has been shown extensively[1, 26, 27] that changes in extracellular stiffness alter the concentrations of nuclear chromatin remodelers, specifically HDAC3 and EZH2 (which are histone methyltransferase/HMT). In this section, we show that our model can be extended to super-resolution imaging experiments published by Heo et al.[1], where chromatin architecture remodeling of a human mesenchymal stem cell (hMSC) is observed in response to changes in microenvironment stiffness. For this analysis, we used hMSC chromosomes 19-22 data-informed polymers constructed from histone ChIP-seq data (covered in methods 2a).

#### Chromatin domains decondense on a stiffer substrate in hMSCs

Heo et al. (2020) cultured human mesenchymal stem cells (hMSCs) on methacrylated hyaluronic acid (MeHA) hydrogels with distinct stiffnesses (30 kPa and 3 kPa) for 48 hours (Fig. 6a). STORM imaging with H2b histone protein labeling revealed a global view of the spatial organization of the genome, revealing distinct chromatin domains with significantly different size distributions between the two samples (Fig. 6b). The softer substrate featured larger domains with more compacted chromatin, whereas the stiffer substrate displayed smaller domains with more decompacted chromatin (more details are provided in SI3.1). A relative reduction of 8% was observed in the domain sizes from soft to stiff substrates. Histone tail mark analysis revealed a substrate stiffness-dependent increase in H3K4me3 and a decrease in H3K27me3, indicating epigenetic regulation of chromatin reorganization. Treatment with the ROCK inhibitor Y-27632 abolished the substrate stiffening effect, confirming that enhanced cellular contractility drives this global increase in nuclear acetylation and chromatin decompaction (data in SI3.2). Consequently, hMSC chromatin reorganizes into a more decompacted state on stiffer substrates owing to altered cellular contractility.

**Figure 6:**
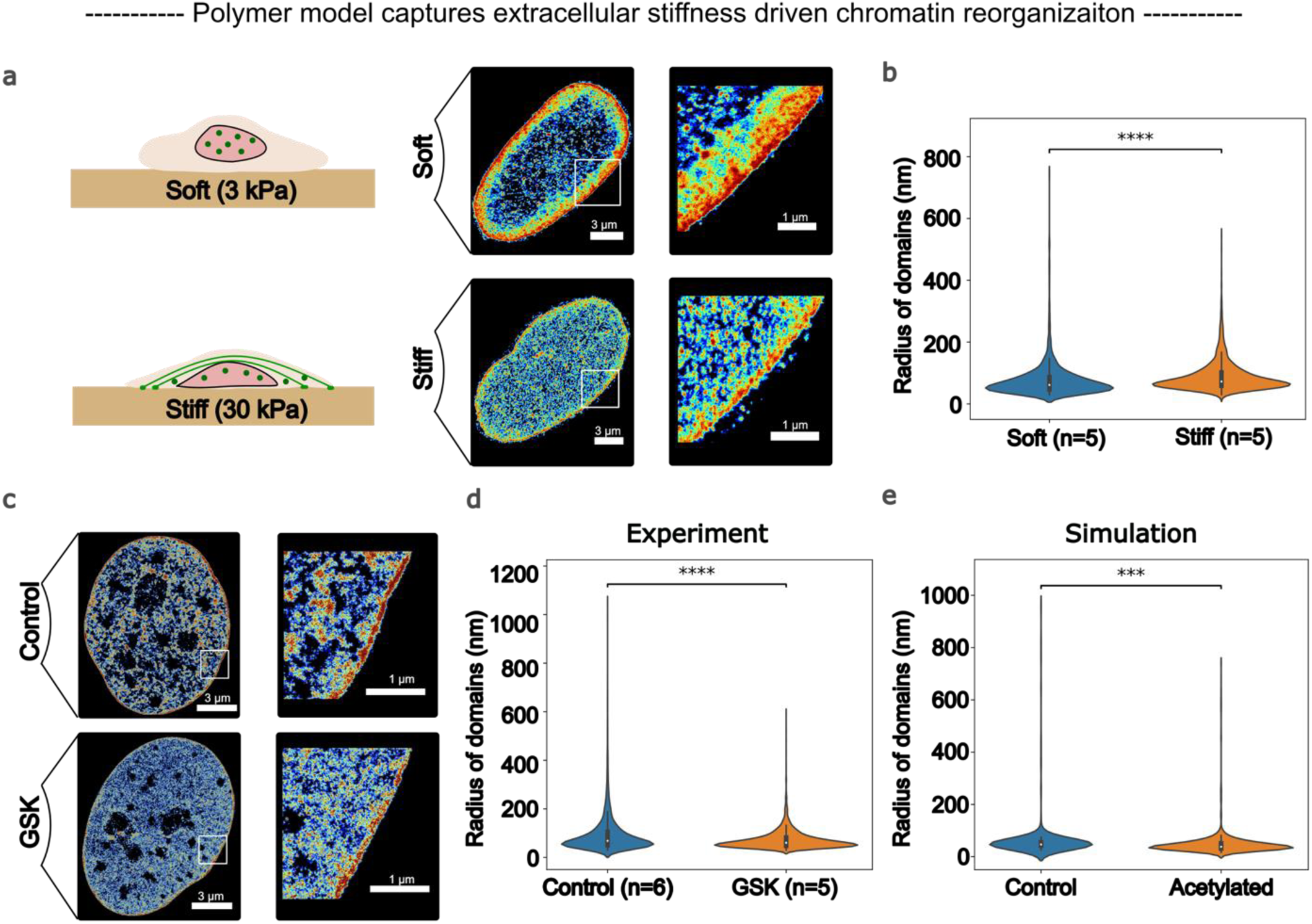
Polymer model captures chromatin reorganization in response to changes in microenvironmental stiffness. a) hMSCs are cultured on 3 kPa (soft) and 30 kPa (stiff) substrates. H2b is labeled and imaged. The chromatin domains decrease in size on a stiffer substrate because of changes in the concentrations of nuclear epigenetic remodelers (represented by green dots) [1]. b) An increase in substrate stiffness drives the decompaction of chromatin, resulting in a decreased observed radius of the chromatin domains. c) GSK treatment of hMSCs inhibits EZH2 (an HMT), leading to lower methylation levels and thus chromatin decompaction, as shown in the STORM images. d) GSK treatment decreases the observed domain sizes, captured through the observed radii of the clutches. e) Simulation of the hMSC data-driven polymers for chromosomes 19-22 replicates the observed decrease in domain size.

#### Chemically induced HMT inhibition leads to chromatin domain decompaction

To ascertain whether H3K27me3 changes can also induce chromatin reorganization, EZH2, a histone methyltransferase crucial for H3K27me3 catalysis, was inhibited by Heo et. al. Using GSK343 (GSK), a specific EZH2 inhibitor, it is confirmed that chromatin structure alterations result from epigenetic profile variations mediated by epigenetic remodelers. Previous studies[1] have shown that on stiffer substrates, EZH2 relocates out of the nucleus, increasing the proportion of euchromatin due to decreased H3K27me3 levels. It is expected that GSK treatment would yield a similar trend as switching to a stiffer substrate. Human mesenchymal stem cells (hMSCs) were treated with GSK throughout their two-day culture period, and subsequent STORM imaging revealed significant chromatin decondensation, characterized by a reduction in condensed chromatin domain size (marked by an 18% reduction in domain size upon treatment) (Fig 6c, d) (more details are provided in SI3.1). This finding confirms that the H3K27me3 changes observed with changing substrate stiffness are sufficient to drive chromatin reorganization. Thus, this presents an ideal scenario for the application of our model, where microenvironmental cues govern chromatin remodeling through nuclear epigenetic tuning.

#### The polymer model captures changes in chromatin organization in response to mechanical cues, specifically stiffness

We validated our model predictions by comparing them to experimentally observed chromatin reorganization trends. Since changes in the concentrations of nuclear epigenetic remodelers, specifically EZH2, drive chromatin reorganization in response to both stiffness changes and HMT inhibition (GSK treatment), we simulated this phenomenon by adjusting the epigenetic reaction rates, which capture the effect of EZH2 in our model. Control simulations were run with a constant net epigenetic makeup, whereas simulations with decreased methylation levels (representing increased stiffness and GSK treatment) were run with adjusted acetylation-to-methylation reaction rates (0.2). The resulting domain size distributions (Fig. 6e) strongly resemble the experimental observations, with a simulated reduction in the mean domain radii of 15%, which aligns with the experimental trends.

Finally, our polymer model can be integrated with our previously published models, linking actin polymerization to nuclear epigenetic remodeler concentrations[26, 27]. This hybrid framework enables the prediction of chromatin organization changes in response to alterations in the cellular cytoskeletal network across diverse microenvironments. By combining epigenetic marks and chromatin organization changes, our model captures the genomic effect of mechanical alterations in tissue properties that occur in diseased states, such as tendinosis. Notably, our model reveals that domain size distribution changes in tendinosis following the chromatin reorganization principles we revealed, replicating the observed chromatin domain scaling trends in cases where nuclear histone epigenetic remodelers play a key role (data in SI3.2)[1]. This phenomenon is particularly significant in response to changes in microenvironmental stiffness, highlighting the model’s potential to elucidate the complex interplay between mechanical and epigenetic regulation in disease.

### 3f Polymer model uncovers a robust mechanism of microenvironment memory formation

Epigenetic memory becomes particularly relevant when the cell passes through multiple microenvironments and has the potential to experience lasting influences from these environments. This is important for processes such as metastasis, wound healing, or morphogenesis, where the cells go through rapidly changing environments, raising the question of how and where the memory of the microenvironment is being stored. Recent work[55] has shown how 3D organization, along with a limitation in epigenetic modifying enzymes, can lead to efficient passage of epigenetic memory from one generation to the next. Our model provides an opportunity to examine how exposure to an environment affects the cell phenotype during the cell cycle. To address this aspect through our model, we take the example of chromosome 19 in the hMSC cell line, which is mesenchymal in nature, with a plastic phenotype compared to fully differentiated cells. To understand the long-term behavior in a varying microenvironment, we start by taking the active-repressed compartmentalization as dictated by the ChIP-seq data (section 2a). We then increased the acetylation for time t’, followed by an increase in methylation for time t’, amounting to a total simulation of 2t’. In an experimental setting, this would correspond to the stiffening and subsequent softening of a substrate (with changes in nuclear HDAC, as shown in section 3b) or to TSA treatment with subsequent washout. In the plotted simulation, the time t’ is chosen to be ∼5 hours of real time, and the acetylation-to-methylation ratio is chosen to be 0.8 for t’ and reduced thereafter, as depicted in the kymograph in Fig. 7a. We observe that the domain sizes decrease with acetylation, followed by an increase with increased methylation (Fig. 7b).

**Figure 7:**
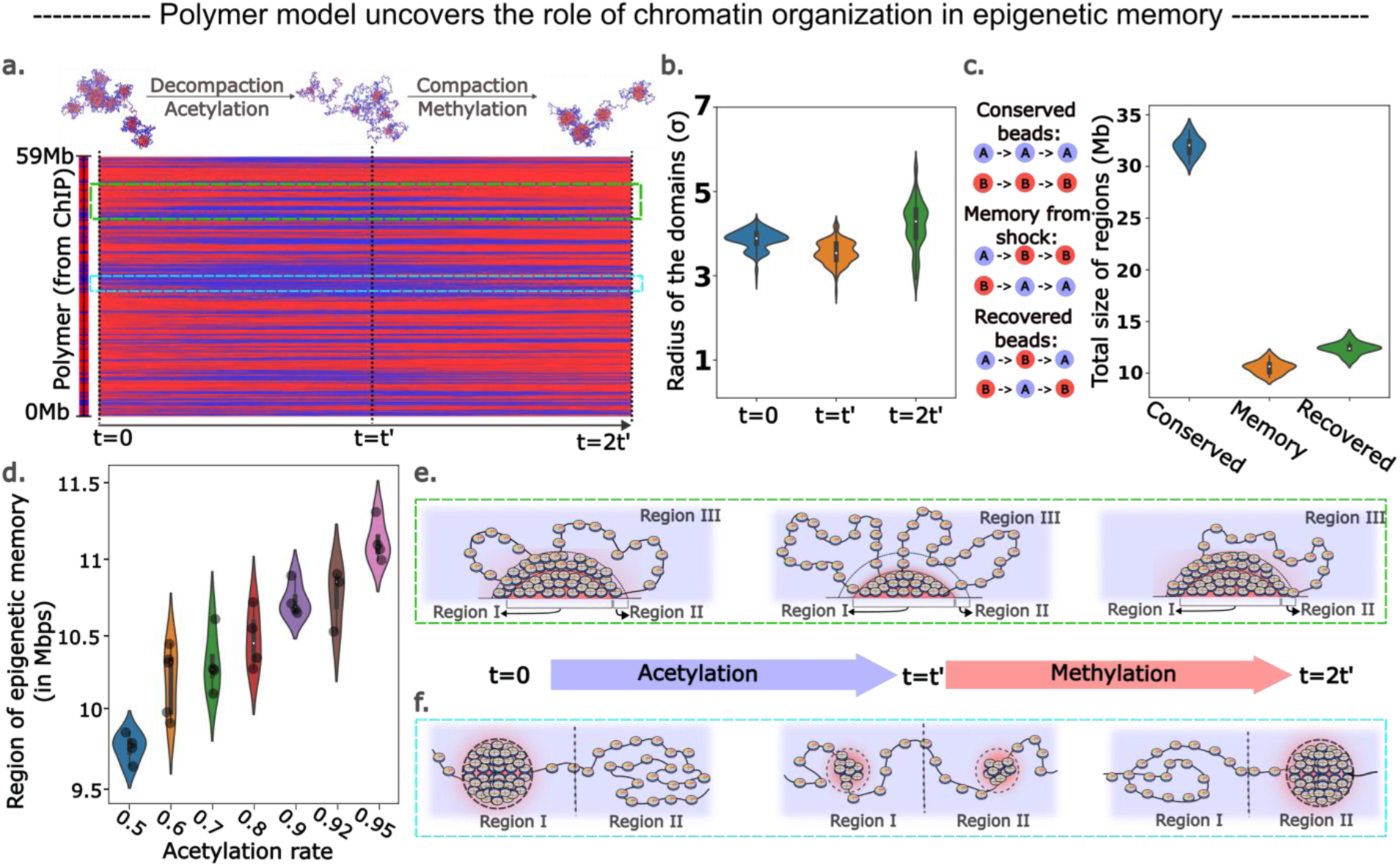
Polymer model reveals a robust mechanism for chromatin organization-driven epigenetic memory. a) Initial acetylation of chromosome 19 from hMSCs (from t=0 to t=t’) followed by methylation (t=t’ to t=2t’). This process is accompanied by simultaneous chromatin decompaction and then compaction. b) Change in the chromatin radius upon acetylation and subsequent methylation. The domain sizes first decrease in size because of decompaction resulting from acetylation and then grow because of compaction due to methylation. c) Evolution of the chromatin is shown through three types of beads: conserved, memory and recovered. Conserved beads do not change their epigenetic marks throughout the simulation time. Beads belonging to the memory from shock category switch their epigenetic mark during acetylation but do not recover during methylation. Recovered beads lose their original epigenetic mark during acetylation but regain it during methylation. Their total size on chr19 is plotted based on multiple final polymer configurations. d) Dependence of the total memory bead segment is plotted as a function of the extent of acetylation. Higher acetylation leads to higher memory of the intermediate state in the same time frame. e) Schematic showing the behavior of large heterochromatin domains through the simulation. The corresponding simulation region is shown in the green box in subfigure (a). The beads in regions I and III are largely conserved, and their epigenetic marks are maintained throughout the simulation. Beads in region II contribute to recovered beads and memory beads, switching their epigenetic marks during acetylation but regaining their original marks or switching and not coming back. (f) Schematic showing the behavior of smaller heterochromatin domains through the simulation. The corresponding simulation region is shown in the cyan box in subfigure (a). Region I loses its epigenetic marks during acetylation and decreases in size. Another neighboring region, region II, forms a nucleating heterochromatin domain. As the methylation rates increase, region II grows to form a distinct domain.

To analyze the memory-like behavior in our system, we look at 3 different kinds of polymer beads, according to their epigenetic trajectory (Fig. 7c): (i) we refer to beads that never change their marks as “conserved beads,” (ii) the beads that change their epigenetic character upon exposure to acetylation (stimuli) and never switch back are referred to as “memory from the shock” and (iii) “recovered” beads are those that change their epigenetic mark but return to their original epigenetic mark after the stimulus is removed. These three types of regions on the chromatin thread highlight the major behaviors one would expect to see in the cell when undergoing extracellular: (i) some phenotypes that are “conserved,” (ii) another set of phenotypes that switch permanently based on external cues and, finally, (iii) a set of phenotypes that change elastically with the microenvironment. We observe that the extent of these regions is sensitive to the magnitude of extracellular stimulus, with more irreversible changes (higher proportion of memory beads) as the external cues increase (Fig. 7d). Detailed kymographs are plotted in SI2.3.

In addition to the central role of the sensitivity of the domain boundaries in response to external cues, we find that regions flanking the domain boundaries play a central role in determining the eventual epigenetic and therefore transcriptional state of the chromatin. In this context, we subcategorize the domains into large domains and smaller domains (as shown through the green and cyan panels of Fig. 7a) to help us understand their evolution better. For the larger domain, we show the corresponding evolution in Fig. 7e. We have three different regions, Regions I, II and III. Both Regions I and III, which are spatially far from the domain boundaries, maintain their epigenetic and therefore accessibility characteristics. Region II, which is at the boundary, undergoes an epigenetic and accessibility cycle throughout the simulation. Through these changes, this region does not recover fully to its original state and contributes to both the formation of permanent epigenetic memory and epigenetic recovery. In contrast, all regions of smaller domains (cyan panels in Fig. 7a and 7f), which are spatially in the vicinity of domain boundaries, are destroyed by shifts in the epigenetic makeup (Fig. 7f, Region I) and new domains are formed through a nucleation and growth process (Fig. 7f, Region II). Therefore, smaller domains can act as domain boundaries themselves and give rise to lasting memory characteristics.

Through this analysis, we hypothesize the central role of domain boundaries in the establishment, maintenance, and dissipation of extracellular memory. Since we observe interesting plastic dynamics even with a coarse-grained system and two epigenetic flavors; in a more realistic system, multiple epigenetic states and their differential propensity to switch can give rise to highly complex cellular phenotype evolution dynamics.

## 4 Discussion

Over the past decade, revolutionary advances in experimental techniques, particularly super-resolution imaging (STORM[95], PALM[96], ChromSTEM[97], etc.) and next-generation sequencing (Hi-C[8], ChIP-seq[16], ATAC-seq[98], etc.), have transformed our understanding of the three-dimensional organization of the mammalian genome and its role in determining the transcriptional state of the cell. Concurrently, significant strides in the analytical and computational modeling of chromatin[99–102] have established a biophysical framework for understanding chromatin packing mechanisms and the pivotal role of epigenetic markers and looping in driving this organization. These breakthroughs have elucidated the multiscale structure of chromatin, spanning from chromatin packing domains[23], visualized through super-resolution imaging, to compartments, topologically associated domains (TADs), and loops, which have been brought to light, primarily through chromatin conformation capture mapping techniques[103]. Data-intensive molecular dynamics approaches[104] and subsequent deep learning efforts[105] have successfully predicted chromatin capture maps via DNA sequences and epigenetic tracks as inputs, demonstrating a strong correlation of compartmentalization with epigenetic marks and their associated proteins. Furthermore, integrated experimental-modeling approaches have provided insight into the dynamic reorganization of chromatin during the transition from interphase to mitosis[106], revealing the crucial roles of looping motors[49] and transcription-driven supercoiling dynamics in chromatin compaction and reconfiguration[14, 107, 108].

Despite significant progress in understanding the relationships between chromatin organization and the epigenetic machinery, the effects of far-from-equilibrium epigenetic remodeling processes on chromatin structure and function remain comparatively less understood. Among the various contributing factors to out-of-equilibrium chromatin dynamics, cohesin-and condensin-dependent looping mechanisms are very well understood[49]. The fact that cohesin depletion leads to a noisier expression pattern rather than significantly altered gene expression suggests the importance of other mechanisms at play[109]. In this light, the role of epigenetic-reaction-dependent chromatin domains, which persist even after Rad21 depletion[110], prominently observed through super-resolution imaging, is worth exploring. Therefore, in the current work, we aimed to explore the role of histone epigenetic remodelers (e.g., HDACs and HMTs) on these chromatin domains in dictating chromatin organization. Efforts to understand other epigenetic remodeling mechanisms, such as the spreading of epigenetic marks across cell replication cycles[45, 54, 55], have been carried out, but to the best of our knowledge, no molecular dynamics-based polymer model has explicitly considered diffusion and microenvironment-driven epigenetic reactions to explain chromatin organization through the formation of heterochromatin-rich packing domains. Crucially, current models are not able to explain the ability the methylation and acetylation reactions in the formation and size scaling of chromatin domains in different scenarios, including disease, development, aging, and the cellular response to drugs. In this study, we take a hybrid approach, integrating simulations with sequencing and imaging experiments to address this knowledge gap. Our molecular dynamics-based model has the following key distinctive features:

1. We employ a data-driven methodology, utilizing next-generation sequencing to inform epigenetic labeling of the chromatin polymer. This enables the design of *in silico* experiments that can be directly compared with alterations in chromatin architecture resulting from changes due to extracellular cues, ranging from mechanical cues, such as substrate stiffness, to the impact of pharmacological and epigenetic drugs. This facilitates a robust validation framework to test the model.
2. Our model incorporates the impact of changes in histone methylation and acetylation on chromatin organization by modeling histone epigenetic remodelers through epigenetic reactions. This feature enables the simulation and analysis of chromatin architectural changes in response to various stimuli within a unified framework.
3. Through chromosome-scale simulations, our model bridges scales by capturing chromatin structure at the nanoscale while simultaneously tracking alterations across multiple kilobases. This allows us to integrate STORM imaging observations with local perturbations detected by next-generation sequencing, providing a unified understanding of chromatin organization.

In our far-from-equilibrium chromatin dynamics model, euchromatin and heterochromatin segments are assigned via binarized ChIP-seq or Hi-C data. To account for the interplay between epigenetics and the chromatin configuration, our model accounts for energetically driven chromatin‒chromatin interactions leading to phase separation of epigenetic domains through Brownian motion and epigenetic diffusion. The out-of-equilibrium dynamics are introduced through epigenetic reactions that capture the action of histone epigenetic remodelers and control the active interconversion of heterochromatin and euchromatin states. Recent work[1, 26, 27] has shown that these epigenetic remodelers are highly sensitive to the microenvironment. Hence, through the current model, we predict the resulting epigenetic landscape and chromatin accessibility, given the initial epigenetics and the variations in the extracellular-driven nuclear concentrations of histone epigenetic modifiers.

Using this epigenetic reaction-driven approach, we demonstrate that the proposed dynamics capture the formation of characteristic nanoscale heterochromatin-rich domains. Our model reveals that heterochromatin-rich packing domain formation results from competition between passive diffusion and active epigenetic reactions, wherein the passive component drives the system toward Ostwald ripening, whereas the active reactions oppose this through energetically unfavorable interchange of epigenetic marks. This competition leads to the scaling of the domain radii with 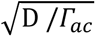, where D is the diffusion constant and *Γ*_*ac*_ is the acetylation reaction rate. We have also shown that the radius of chromatin domains changes nonlinearly with changes in epigenetic reactions, with the radii of the domains scaling proportional to 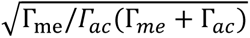, capturing the changes in the chromatin organization with changes in the histone epigenetic remodeler concentrations. We confirmed this size-scaling behavior of the chromatin domains through super-resolution imaging of unperturbed and hyperacetylated A375 melanoma cells via TSA (HDACi) treatment.

Next, we utilized our polymer model to investigate kilobase-scale changes in response to changes in epigenetic reaction rates. Our model reveals that, at the single-cell level, while changes in epigenetic reactions lead to genome-wide epigenetic and conformational perturbations, they are more likely to occur at the kilobase-scale boundaries of heterochromatin domains. When observed over a population of cells, analogous to bulk sequencing of heterogeneous cell populations, our simulations show that these epigenetic shifts progressively concentrate at domain boundaries. These findings suggest that population-level sequencing methodologies such as Hi-C and ChIP-seq are likely to overlook cell-to-cell heterogeneity, which single-cell techniques can capture. We find that imaging techniques, and in general single-cell techniques, can reveal drug exposure changes that are not robust across populations. While on the contrary, bulk sequencing technologies excel at obtaining dominant population-wide changes. To validate our epigenetic switching predictions, we performed Hi-C sequencing on hyperacetylated melanoma cells and observed concentrated epigenetic shifts at domain boundaries without widespread compartmental changes. RNA-seq analysis revealed a correlation between epigenetic landscape alterations and transcriptional state changes in melanoma cells, with boundary region genes following local epigenetic and accessibility trends. Notably, key EMT-related genes near domain boundaries were upregulated, highlighting the crucial role of domain boundaries in determining cell fate. Our predictions were further validated by super-resolution imaging of hMSCs on substrates of different stiffnesses, demonstrating the versatility of our approach in capturing chromatin organization changes. Our *in silico* experiments demonstrated how microenvironment-induced changes in chromatin modifier levels can lead to genome structure rearrangement and cell phenotype changes and revealed that epigenetic memory is predominantly encoded in regions flanking domain boundaries. The extent to which these flanking regions are affected by epigenetic reactions is variable and increases with the magnitude of the shifts in the magnitude of the extracellular stimulus. Ultimately, our hybrid experimental-simulation approach elucidates the pivotal role of domain boundaries in maintaining cellular identity.

In summary, our chromatin polymer model yields two key findings, which we distill in our schematic (Fig. 8):

1. Chromatin domain formation and scaling emerge from the interplay between passive diffusion and active epigenetic reactions, as validated by super-resolution imaging of A375 melanoma cells. Furthermore, we revealed that chromatin domains exhibit scalable responses to substrate stiffness, adhering to the same scaling laws.
2. Changes in epigenetic reactions primarily impact the epigenetic landscape at domain boundaries, driving transcriptional state changes, as validated by Hi-C sequencing and RNA-seq. As a result, domain boundaries can also serve as critical control regions for the formation of microenvironment-driven epigenetic memory.

**Figure 8:**
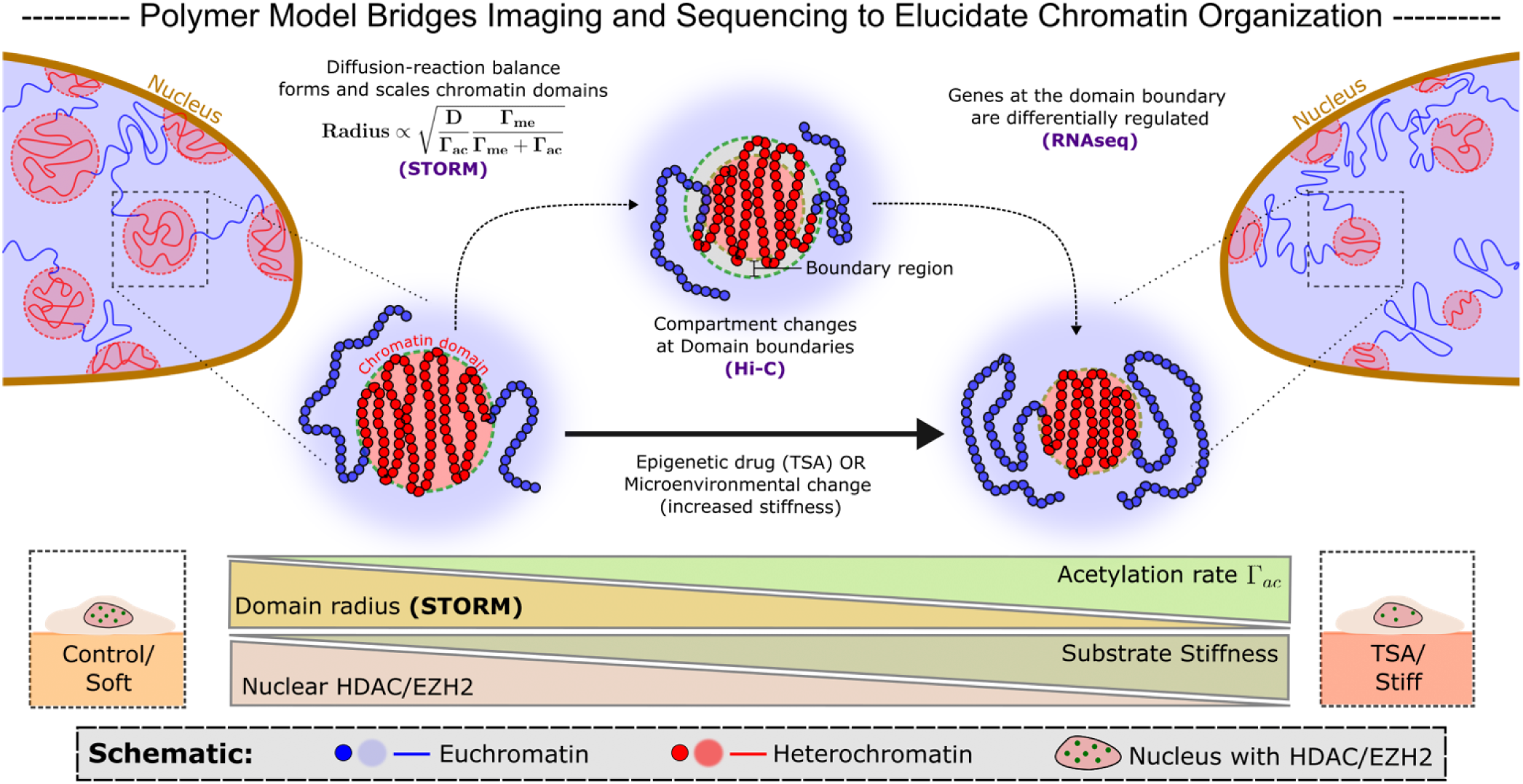
Polymer model bridges STORM, Hi-C and RNA-seq to reveal the importance of domain driven chromatin organization and domain boundaries in genome regulation. Our model shows that heterochromatin-rich chromatin domains form through a balance between diffusion and epigenetic reactions. Additionally, a scaling relation between the domain size distributions and the epigenetic reaction rates is established which capture the evolution of the chromatin organization with changing histone epigenetic remodeler concentrations. These scaling relations are validated through STORM imaging of A375 cells with TSA treatment and hMSCs with substrate stiffness change and GSK treatment. The model also predicts that epigenetic changes occur at the domain boundaries prominently. These predictions are confirmed by Hi-C sequencing with a high accuracy. The epigenetic changes lead to transcriptional changes close to the domain boundaries, especially affecting genes which are important for the cell line. In the case of A375 melanoma cells, we showed EMT-specific genes around the domain boundaries being affected. Lastly, we show that epigenetic memory can also stored around the domain boundaries.

Our study provides a comprehensive understanding of chromatin dynamics and epigenetic regulation, underscoring the pivotal role of domain boundaries in shaping cellular identity. By demonstrating the applicability of our model to various scenarios where histone read-write mechanisms govern epigenetic regulation, we reveal the far-reaching implications of our findings. Notably, our analysis of Y-27 treatment and tendinosis highlights the model’s utility in understanding biochemical treatment and disease progression. Moreover, the importance of histone epigenetic alterations extends to various diseases, including cancer onset and metastasis[111], Alzheimer’s disease[112], and cell fate decisions[113]. Crucially, our work has the potential to reveal the central role of histone epigenetic modifiers, which can pave the way for discovery of pharmacological interventions against cancer[114] and diabetes[115], among other disorders.

The formation of nanoscale chromatin domains creates a rich heterochromatin‒euchromatin interface at boundaries, which are energetically more susceptible to changes due to epigenetic enzyme fluctuations than other loci. Recent studies using dCas9-KRAB constructs support this notion[116], highlighting the dynamic nature of these boundaries. The recruitment of the RNAPII transcriptional machinery to these interfaces[107] further underscores its critical role in regulating the cell’s epigenetic landscape. We propose that domain boundaries can serve as key chemo-mechanosensors, translating external cues into epigenetic changes that determine single-cell fate. This cell-invariant mechanism can enable cell line- and cell history-specific responses, which integrate with established signaling pathways to orchestrate transcriptional programs. For example, substrate stiffness-mediated activation of mechanosensitive pathways such as YAP/TAZ in mesenchymal cells[117] are well established, but our model demonstrates that a cell’s microenvironmental history, encoded in its epigenetic and spatial configuration, can influence its response to such stimuli, yielding diverse outcomes. Our findings underscore the potential role of epigenetic landscape-driven domain boundary positioning in shaping cell-specific transcriptional programs.

While our model has proposed key biophysical mechanisms governing chromatin organization and gene regulation, future refinements can address current limitations. Notably, incorporating multistate chromatin models that account for different epigenetic flavors (such as ChromHMM[2] or MEGABASE[35]), chromatin‒lamina interactions[118], and sequence-dependent epigenetic marking[119] will increase the predictive power. Integrating replication dynamics and epigenetic spreading mechanisms[120] will reveal long-term chromatin architecture changes, which are crucial for understanding cancer metastasis and development. Furthermore, exploring the interplay between chromatin organization and nuclear signaling pathways will provide a more comprehensive understanding of cell fate determination. By building upon our foundation, future studies can investigate the role of chromatin organization in disease mechanisms, ultimately informing novel therapeutic strategies. The integration of experimental and theoretical approaches will be essential for achieving a holistic understanding of chromatin organization and its role in governing cell fate.

## Acknowledgments

The authors thank Aayush Kant, Monika Dhankhar and Zixian Guo for the multiple useful conversations, suggestions and for sharing their work on the STORM analyses code. We thank Christopher Playter and Priyojit Das (UTK) for assisting with the Hi-C experiments and analyses. This work was supported by NIH Award U54CA261694 (V.B.S.); NCI Awards R01CA232256 (V.B.S.); NSF CEMB Grant CMMI-154857 (V.B.S.); NSF Grants MRSEC/DMR-1720530 and DMS-1953572 (V.B.S.); and NIBIB Awards R01EB017753 and R01EB030876 (V.B.S.). This work was supported in part by the National Institutes of Health [NIGMS grant R35GM133557 to R.P.M].

## Author contributions

V.V. and V.B.S. conceived the polymer model. V.V., R.B., and L.S. carried out the computational and numerical analyses. R.G. and R.P.M carried out the sequencing experiments. V.V., R.G., and R.P.M carried out the sequencing analyses. V.V., J.T. and M.L. carried out the super-resolution imaging and analyses. All authors contributed to writing the draft.

## Data availability

The data supporting the findings of this study are available from the corresponding author upon request. The data generated in this study are provided in the Source Data file. The sequencing raw files are available on the GEO dataset with the accession number: GSE275755.

## Code availability

The code used for measurement of sizes of heterochromatin domain obtained from STORM imaging is freely available through GitHub (https://github.com/ShenoyLab/STORM_Analysis_Parameter_Extraction). The simulations have been done in LAMMPS (stable release Aug 2023) with custom fixes written while the analyses is done in python. Both these are available at: https://github.com/vinayakv161/Epigenetic_diffusion_and_reactions. The Hi-C analyses is performed using cooltools[121] and RNA-seq using DESeq2[72] and clusterprofiler[74].

