## Supplementary Information for "Polymer Model Integrates Super-Resolution Imaging and Epigenomic Sequencing to Elucidate the Role of Epigenetic Reactions in Shaping 4D Chromatin Organization"

[1] Center for Engineering Mechanobiology, University of Pennsylvania, Philadelphia, PA, 19104
[2] Department of Materials Science and Engineering, University of Pennsylvania, Philadelphia, PA, 19104
[3] Department of Physiology, Perelman School of Medicine, University of Pennsylvania, Philadelphia, PA 19104, USA
[4] Department of Biochemistry & Cellular and Molecular Biology, University of Tennessee, Knoxville, TN 37996, USA
[5] Department of Bioengineering, Stanford University, Stanford, CA 94305, USA
* Corresponding author

**Contents:**

**1. Details of the polymer model**
- **1.1. Building the chromatin polymer from sequencing data**
- **1.2. Potentials used in the polymer model**
- **1.3. Details of the executed dynamics**
- **1.4. Detailed balance considerations**
- **1.5. Running simulations and computing observable quantities**
- **1.6. Analytical derivation of domain scaling**

**2. Cell-line specific simulations on data-informed polymers**
- **2.1. Domain stability for other chromosomes of A375 and hMSC**
- **2.2. Altered diffusion, acetylation and methylation runs**
- **2.3. Memory analysis**

**3. STORM analysis**
- **3.1. Imaging analysis workflow for calculating domain sizes**
- **3.2. Data for domain size distribution of Y-27 treatment and tendinosis**

**4. Sequencing analysis**
- **4.1. Analyzing the HiC map for A375 cells**
- **4.2. RNA-seq analysis for A375 cells**

### 1 Details of the polymer model

#### 1.1 Building the chromatin polymer

The polymer model presented in this study leverages experimental data to establish the initial configuration of the chromatin polymer. We utilize either ChIPseq in the form of Broad ChromHMM[2] tracks or experimentally obtained HiC data to determine the bead labels as euchromatin-like/active (blue) or heterochromatin-like/repressed (red). For the chromosomes of IMR90 and hMSC cell lines, we employ publicly accessible ChromHMM data for polymer initialization[3]. Specifically, we utilize the 15-state ChromHMM information and assign states 1-11 as blue beads corresponding to active states, while the remaining states are designated to red beads corresponding to repressed states. This strategy has been adapted from the CCM model[4]. Given that our bead size (10 kbp) is significantly larger than the 200 bp resolution of ChromHMM, we use a majority rule approach, where each bead is assigned to the state that occurs most frequently within its boundaries. Although this coarse-graining approach results in some loss of information, we observe that these regions predominantly belong to a single state, ensuring that the polymer faithfully represents the underlying epigenetic structure. For A375 cells, we utilize experimentally obtained HiC of the cells in control condition to construct the initial polymer. Here, we compute the first eigenvector (PC1) from the experimentally derived HiC contact map using cooltools[5] to delineate the A and B compartments. Beads falling within the A compartment are assigned as blue, while those falling within the B compartment are designated as red. More information on the Hi-C analysis is provided in section 4.1. All the polymer strands we form in the study are shown in figure Fig. S1. Moreover, the sourcing and reason to use the different cells is depicted in Table S1.

For our simulations, we chose a bead size $(\sigma)$ of 65 nm corresponding to 10 kbp of contiguous chromatin region. A 10 kb region would dictate the number of nucleosomes of the order of ~50 contained in the bead. Using a lower limit, closed packing case, we can equate the volume of the bead to the volume of 50 nucleosomes, which would give us a lower bound of $\sigma =$ 40nm, assuming the size of nucleosomes to be ~10nm. For the upper bound, we can consider the polymer as a worm-like chain in which case we can get the size using the persistence and contour lengths as $\sigma^2 = 2L_C L_P - 2L_P^2$(assuming persistence length is significantly smaller than contour length, which is true for our case)[6]. Assuming a persistence length of ~1000 bp[7], we get $L_P = 6(16.5 + 10)$nm (with 16.5nm being the length of linker DNA and 10nm as the nucleosome size)[4]. This gives us an upper bound of ~500nm. Based on these estimates the choice we make here is in a physically reasonable range, if not a bit conservative. The conservative choice is reinforced by experimental observations made in Boettiger et. al.[8] where our bead size fits perfectly in scaling discovered. Moreover, another recent work[9] has shown that a 5kb region is well represented by a bead of size ~80 nm which falls in a similar range. To maintain a packing density consistent with experimentally observed ranges, the polymer is positioned within a fixed wall box, targeting a biologically relevant volume fraction of ~0.15[10, 11].

To understand the general physical principles driving the formation and growth of chromatin clutch domains, we also use a toy polymer model to elucidate the physical underpinnings of the proposed model. For this purpose, we choose a 3000-bead (~30Mbps) polymer and randomly assign the bead labels to repressed (active) marks with a probability of 0.6 (0.4). This ratio has been chosen to account for the repressed majority (including the centromeric and telomeric regions) found in all the chromosomes we considered for the study.

In this work, we do not explicitly consider the role of loop formation through cohesin mediated loops. Even though they form a central part of chromatin organizational structure, we do not consider their role due to the following two reasons: a) Our current resolution is at 10kbps per bead and an average CTCF based cohesin loop is about ~50-100kbps. Hence it would be a loop only over 5-10 beads. b) CTCF ChIP-seq is not available for all the cell lines used and hence to keep our observations consistent, we did not use CTCF mediated loops for the cell lines we chose. We do not though, that our model does allow simple introduction of the loops in a static or dynamic manner. For the IMR90 cells, we ran simulations with static CTCF mediated loops which were added in a manner like Shi et. al. [4] and found that the obtained changes in epigenetic landscape did not show any differences from the case where such loops were not included. We do acknowledge that at a finer scale, the role of such loops will become more important as they can govern the local neighborhood of a chromatin segment, which ultimately determines the epigenetic fate of that segment.

##### 1.2 Potentials used in the polymer model

We use the LAMMPS[12] engine to simulate our molecular dynamics system. The chromatin is modeled as a copolymer with euchromatin/A compartment/active regions represented as blue beads and heterochromatin/B compartment/repressed regions represented as red beads. The interaction between these beads is enforced using a Lennard-Jones potential as follows:

$$U_{LJ}^{\alpha\beta} = \begin{cases} 4\epsilon_{\alpha\beta}\left[\left(\frac{\sigma}{r}\right)^{12} - \left(\frac{\sigma}{r}\right)^{6}\right] & for \quad r \le r_{cut} \\ 0 & for \quad r > r_{cut} \end{cases}$$

where $\alpha$ and $\beta$ can be blue or red based on the ChIPseq/HiC annotation. The bead $\sigma = 1$ is chosen to be the same for both the beads. The radius for blue beads should be higher since the chromatin compaction is less but we keep it this way to make reduce model complexity. The interactions between red and blue beads are set such that the potential cutoff distance $r_{cut}$ is set to $2^{1/6}\sigma$ resulting in excluded volume interactions also known as the WCA potential. The use of this potential is granted given the short-range interactions with respect to the length scale we have chosen for the system. The intra-cutoff $r_{cut}$ among A or B beads is set at $1.8\sigma$ to account for bridging protein driven self-attraction. This leaves us with tuning the interaction energies between the heterochromatin-heterochromatin beads ($\epsilon_{HH}$) and the euchromatin-euchromatin beads ($\epsilon_{EE}$). Since heterochromatin is more compact that euchromatin, we add an additional constraint that $\epsilon_{EE} < \epsilon_{HH}$. In addition to this we have FENE (Finite Extensible Non-linear Elastic) potential mediated bonds in between neighboring beads on the linear polymer to maintain the integrity of the polymer:

$$U_{FENE}^{i} = -\frac{1}{2}K_{FENE}R_0^2 ln\left[1 - \left(\frac{|\boldsymbol{r}_{i+1} - \boldsymbol{r}_i|}{R_0}\right)^2\right]$$

where $K_{FENE}$ is the spring constant and $R_0$ is the equilibrium bond length.

Combining the two, the net potential attributed to any bead $i$ of the system can be described as the following function:

$$U_{net}^{i}(x) = \frac{1}{2}\sum_{j \neq i} U_{LJ}^{ij} + \sum_{j \in \{i-1, i+1\}} U_{FENE}^{j},$$

where $i$ covers all the beads in the system. Since the persistence length of the chromatin fiber has been established to be about ~1000 bp[7] and our beads are much bigger than that, we ignore any polymer stiffness apart from the stiffness provided by excluded interactions between the beads.

To obtain the initial configuration of the polymer, we form a rod-like initial polymer using the process described in section SI1.1. This polymer is then arranged in a self-avoiding random walk. The Lennard-Jones potentials are turned on and the polymer is relaxed for $10^7$ time steps to form the initial configuration. The potential energy plots, averaged over simulations show that the equilibration is reached (Fig. S2). Since we have only two parameters to tune ($\epsilon_{HH}$ and $\epsilon_{EE}$), with the constraint $\epsilon_{EE} < \epsilon_{HH}$, we use a hit-and-try methodology to analyze how well the obtained mean spatial distance vs. the genomic distance plots agree with the experimentally observed data[13] for IMR90 cells for chromosomes 20 and 21. Among the tested values we found $\epsilon_{EE} = 0.3$ and $\epsilon_{HH} = 1$ fit the experimentally observed values the best as shown in Fig S3. The polymer configuration is further validated through the Hi-C map presented in Fig 1e of the main text. To compute the Hi-C map, multiple polymer relaxations were executed, and any two beads were considered in contact if they were within 0.2 $\sigma$ of each other. Thereafter the contact probability was calculated as a ratio of the number of simulations which maintained the contact. To match it with experimental observations[14], the experimental Hi-C data was normalized and changes to a log-scale.

##### 1.3 Details on diffusion and reactions

We use Brownian Dynamics (shown in Fig S4a) to evolve the system in time in the presence of an implicit solvent, which in our case is nucleoplasm (water):

$$m\frac{d^2\boldsymbol{r}_i}{dt^2} = -\gamma\frac{d\boldsymbol{r}_i}{dt} - \nabla U_{net}^{i}(x) + \boldsymbol{\xi}_i$$

where $\gamma$ is the friction coefficient and $\boldsymbol{\xi}_i$ is a stochastic noise obeying the fluctuation-dissipation relationship $\langle \xi_{i,\alpha}(t)\xi_{j,\beta}(t')\rangle = 2\gamma k_B T \delta_{i,j}\delta(t-t')\delta_{\alpha,\beta}$. Indices $i,j$ run over particles and $\alpha,\beta$ run over the cartesian space. In this setting, using the Einstein relation, we obtain:

$$D = \frac{k_B T}{\gamma} = \frac{k_B T}{3\pi\eta\sigma},$$

where $\eta$ is the viscosity of the background. Considering a bead size of 65 nm for 10 kbp region and a viscosity of $\eta = 150cP$ we can estimate corresponding Brownian time:

$$\tau_{Br} = \frac{\sigma^2}{D} = \frac{3\pi\eta\sigma^3}{k_B T} \approx 0.3\ s.$$

We utilize a velocity Verlet time integration methodology implemented within the LAMMPS engine in an to execute the trajectories in an NVE ensemble. Considering that we have chosen an integration time $\Delta t = 0.01\tau_{Br}$, we simulate approximately $\sim 1hr$ of real time corresponding to $1 \times 10^6$ integration time steps. This is the conversion have used for all the computations performed in this work.

The epigenetic diffusion is modeled like Kawasaki dynamics studied in Ising spin systems (Fig S4b)[15]. We choose an epigenetically conservative exchange process since diffusion does not alter the ratio of hetero-to-euchromatin but drives the system to a lower energy state through epigenetic redistribution. Each exchange between the epigenetic marks (colors) of the beads is modeled using a Metropolis criterion which is dictated by the acceptance probability of exchange as:

$$p(s \rightarrow s') = \min(1, e^{-\Delta E/k_B T})$$

where $\Delta E$ is the energy difference between the final (after epigenetic exchange) and the initial (before epigenetic exchange) configuration. The temperature $T$ is the same as the Brownian simulation to maintain detailed balance and not introduce any out-of-equilibrium phenomenon. This is explained in detail in the following section.

The epigenetic reactions account for microenvironment-driven methylation ($\Gamma_{me}$) and acetylation ($\Gamma_{ac}$) of the chromatin. They are modeled as a Monte-Carlo processes where any bead can be randomly chosen, and its epigenetic marking can be changed from euchromatin to heterochromatin with a probability equal to $\Gamma_{ac}$ /$\Gamma_{me}$ and vice versa (Fig S4c). This introduces non-equilibrium phenomenon by breaking detailed balance as discussed in the following section.

##### 1.4 Accounting for detailed balance in the simulations

Chromatin dynamics in our model are driven by passive diffusion and active epigenetic reactions. It is central to our finding that the out-of-equilibrium epigenetic remodeling is driven solely by epigenetic reactions. To show our proposed dynamics follow these constrains, here we show how well our model follows equilibrium criteria using detailed balance. We use the Kolmogorov criteria to test detailed balance conditions. According to the criteria, for a system exhibiting detailed balance, the transition rates over any closed loop of states of the system should be independent of the direction in which the loop is traversed. We first account for our choice of the temperature of the Brownian dynamics to be the same as the epigenetic diffusion. As shown in Fig S5(left), we consider a system of four particles which are acted upon by Lennard-Jones interactions as described in the previous sections. The initial and final state are the same, thereby constituting a loop. We only consider the effect of diffusion of epigenetic marks and Brownian motion and quantify the probability of each step using $\exp(-\Delta E/kT_x)$, where $\Delta E$ is the energy required for the process and $T_x$ is the temperature for the Brownian movement or epigenetic diffusion. $T_x$ can be $T_E$ for the case when we are attempting an epigenetic diffusion move and $T_{Br}$

for the case of Brownian diffusion. As can be seen, the clockwise (forward) loop in Fig S5 (left panel, it has the following rate:

$$P_{clock} = \exp\left(-\frac{2\epsilon}{k_B T_E}\right)$$

Contrary to this, the rate of the anti-clockwise (backward) loop is given by:

$$P_{anti-clock} = \exp\left(-\frac{\epsilon}{k_B T_E}\right) \exp\left(-\frac{\epsilon}{k_B T_{Br}}\right)$$

where $T_{Br}$ is the Brownian temperature. Detailed balance holds true only if $P_{clock} = P_{anti-clock}$. Hence for detailed balance to hold, $T_E$ must be the same as $T_{Br}$. A simpler phenomenological argument to understand why this must be true is as follows: In a canonical ensemble picture, the Brownian dynamics are driven by a heat bath at temperature $T_{Br}$ and the epigenetic exchange by a bath at temperature $T_E$. If the two temperatures are not equal, then a constant energy flux is established between the two baths, making it an active system.

Using a similar scheme, we next argue how epigenetic reactions break detailed balance in our model setup. We consider a system with two particles which are initially epigenetically distinct and proceed as shown in Fig S5 (right panel). As is clear from the figure the clockwise rate is given by $P_{clock} = \Gamma_{me}\Gamma_{ac}\exp\left(-\frac{\epsilon}{k_B T_{Br}}\right)$ while the anti-clockwise loop rate is equal to $P_{anti-clock} = \Gamma_{me}\Gamma_{ac}$. Hence, irrespective of the value of the reactions, they will always break detailed balance in the current setup as $P_{clock} \neq P_{anti-clock}$. Hence it is only the reactions which introduce an out-of-equilibrium aspect to our model while the diffusions maintain detailed balance.

##### 1.5 Running simulations and computing observable quantities

With the initial configuration in SI1.2, we decide on the reaction rates as described in the main text and run simulations generally for a 2 hour time point or 10 hour time point, corresponding to $2 \times 10^6$ integration steps or $10 \times 10^6$ integration steps respectively. Simulations are repeated over different initial configurations to establish statistical significance and to capture the effects of cellular heterogeneity. Our analysis consists of both, analysis over single simulations and over multiple simulations based on the context. Broadly we use the following analysis strategies to observe key observables in our systems:

1. <u>Measuring the radius of gyration of the heterochromatin-rich packing domains (Fig S6):</u> To identify the domains, we first isolate the heterochromatin (red) beads from our polymer. Thereafter, we run the DBscan clustering algorithm, available through the scikit-learn package[16] in python. For the implementation, we use an eps=1.2 and the minimum number of particles in a cluster as 10. Then we compute the radius of gyration of the obtained domains. We note here that for all the cases where we plot the distribution of their domains over all obtain clusters except for the case with the prototype polymer.
2. <u>Producing kymographs and distance maps to analyze the spatiotemporal evolution of epigenetic marks:</u> We use simple matplotlib based scatter plots to dhow both the kymographs and distance plots. To produce the kymographs, we do not average over

multiple simulations as the aim is to show single cell scale epigenetic evolution while for distance maps multiple simulations are averaged over.

3. Producing polymer figures: We use VMD[17] to visualize the polymer 3D configurations.

##### 1.6 Analytical derivation of domain scaling

Here we take a nuanced approach wherein we try to show the scaling relation through a simple dimensional analysis argument. As shown in Fig S7, across the boundary of a heterochromatin rich domain, there are two fluxes which are balancing the size of the domain:

1. The diffusion of the heterochromatin into the domain: The flow of heterochromatin into the domain can be quantified as the inward flux times the surface area of the sphere. To calculate the flux, we turn to Fick's law, which dictates that:

$$J = -D\frac{d\phi}{dr} = -D\frac{[concentration]}{[length\ scale]}$$

Where $J$ is the flux, $D$ is the diffusion coefficient and $d\phi/dr$ is the heterochromatin concentration gradient. More precisely, we need to realize the best estimates for the concertation and relevant length scale in the system to arrive at a reasonable empirical relation for the flux. Let us assume that due to the formation of a depletion layer right outside the heterochromatin domain, the heterochromatin content is nil. Therefore, the flux now only depends on the far-field concentration. In this case, the far field concentration will be the average heterochromatin content in the system which is $\frac{\Gamma_{me}}{\Gamma_{me}+\Gamma_{ac}}$. The only relevant length scale in the diffusion problem is the radius of the sphere and hence we can estimate the flux as (ignoring the sign):

$$J = D\frac{1}{R}\frac{\Gamma_{me}}{\Gamma_{me}+\Gamma_{ac}}$$

This will dictate that the net inflow of heterochromatin can be estimated as:

$$Influx\ of\ heterochromatin = 4\pi R^2 \times J = 4\pi RD\frac{\Gamma_{me}}{\Gamma_{me}+\Gamma_{ac}}$$

2. The conversion of heterochromatin to euchromatin within the domain due to acetylation reaction: We firstly assume a uniform concentration of heterochromatin within the formed domain, C. Thereafter, since the reactions occur uniformly over the whole space in our setting, we can estimate the outflux of heterochromatin as:

$$Outflux\ of\ heterochromaitn = Vol.of\ sphere \times rate\ of\ acetylation \times C = \frac{4}{3}\pi R^3\Gamma_{ac}C$$

Equating the outflux with the influx we obtain:

$$R^2 \propto \frac{\mathrm{D}\,\Gamma_{me}}{\Gamma_{ac}(\Gamma_{me}+\Gamma_{ac})}$$

The extended derivation for the domain scaling is provided in the SI of Heo et. al.[18] or Kant et. al.[19].

#### 2. Cell-line specific simulations on data-informed polymers

##### 2.1 Domain stability for other chromosomes of A375 and hMSC

In the scenario where the nuclear concentration of epigenetic/histone remodelers is constant, it is imperative to show that our model can maintain the 3d chromatin organization and epigenetic distribution, which we quantify through the kymograph (representative of the epigenomic stability) and the distance matrix (representative of the spatial stability). In the main text, we showed the kymograph of chromosome 19 of A375 cells for ~2 hours of real time. Here we extend that observation to account for the chromatin accessibility. We show the distance matrix for chromosomes 18-21 of A375 cells for about ~2 hours of real time, over which we have done majority of analysis for our paper in Fig. S8. Thereafter, we show the epigenetic stability of these chromosomes for ~10 hours of real time in Fig S9.

Lastly to ensure that such a trend is observed across cell lines, we repeat this analysis for hMSC cells with the distance matrices in Fig. S10 and kymographs in Fig S11. This analysis ensures that in the case where there are no shifts in the average nuclear epigenomic makeup, the chromatin will sustain its epigenomic and pair-wise spatial identity. Since the epigenomics remain stable through the time, this will also ascertain that the transcriptional activity of the cell also remains stable in the interphase time window.

##### 2.2 Altered Diffusion, Acetylation and Methylation runs

One of the key parameters for our simulations is the ratio of the diffusion to reaction rate, i.e. $D/\Gamma_{ac}$. We have already quantified the evolution of the 3D chromatin organization with respect to this parameter in the main text and explained its origin in SI1.6. Here we analyze the evolution of the epigenetic marks on the chromatin strand of A375 cells as a function of his parameter in Fig. S12. The observations are like observations of the mesoscale observations wherein the epigenetic distribution randomizes completely as the reaction rates increase while we observe almost complete Ostwald ripening as the diffusion rates increase.

With a change in the average nuclear concentration of the epigenetic regulators, the epigenetic makeup of the chromatin strand changes. We already show the chr19 specific changes after acetylation for A375 in the main text. Here we extend those results to show how the changes are replicated in other chromosomes of A375 and hMSCs which are central to our study. For A375, we firstly show how the epigenetic evolution of the epigenetic marks on acetylation and methylation through kymographs in Fig. S13. These figures suggest that major chromatin epigenetic changes are constrained to the domain boundaries. To cement this observation, we plot the distance matrices for acetylation and methylation in Fig. S14. To drive home the concept of localization of epigenetic changes to the domain boundaries we further analyze the methylation of chromosome 19 and acetylation and methylation of chromosome 20 in more detail. Thus this analysis confirms that chromatin domains change epigenetically.

##### 2.3 Memory analysis

For the memory analysis, there are two primary parameters which we could vary: i) The total time and ii) The magnitude of the reaction rates. In our would we focus on the later and keep the total time a constant. We do this mainly since we are bound by the timescale of the interphase of a cell which is ~10 hours and our attempt is to see how the organization changes in that timescale. We perform this analysis for chromosome 20 of hMSC cells. The plots for the varying epigenetic reaction rates are plotted in Fig S15.

Our model can be combined with chromatin replication models and longer timescale observations could be made as well but it lies beyond the scope of the current model.

#### 3. STORM analysis

##### 3.1 Computing domains from STORM images

The images obtained from ONI STORM are first processed using the Nanoimager (ONI) software to perform drift correctio. Thereafter, the images are imported to a custom written code on MATLAB code which we have used in our previous publication. Voronoi-tessellation is used for density quantification and construction of a Voronoi polygon of each localization. Then the density corresponding to each polygon is approximated using the inverse area of the polygons. Based on custom threshold, the nucleus is divided into heterochromatin and euchromatin rich regions and the heterochromatin rich areas are conserved. This threshold value is determined such that > 60 percentile of the density distribution fall into the heterochromatin regime and the threshold is conserved through the different replicates. It is important to note that the density threshold is kept same for a treated sample and its corresponding control. Density-Based Spatial Clustering of Applications with Noise (DBSCAN) is then used to obtain the heterochromatin domains from the underlying point cloud distribution. The radii of these domains are what we refer to as the domains in our imaging analysis.

We illustrate this workflow in Fig. S16. We plot one replicate of each of the control and TSA treated cells and along with that plot the DBSCAN detected interior domains (shown in black) for each of the cell. Our workflow can also detect lamina associated domains, which are excluded from our analysis in this work[1]. The domain size distributions for the interior domains show the Control-vs-TSA trends which have been presented in the main text. Similar analysis has been repeated for cells on stiff and soft gels and GSK treatment.

##### 3.2 Data for domain size distribution of Y-27 treatment and tendinosis

The data for Y-27 and tendinosis is obtained from Heo et. al. [18] while the analysis is followed from DGK et. al. [1] and plotted in Fig. S17.

#### 4. Sequencing analysis

##### 4.1 Analyzing the HiC map and corresponding simulations for A375 cells

To obtain the A/B compartments, the two replicates for the control and TSA treated cells were combined to get a more robust Hi-C signal. Thereafter principle component analysis was performed using cooltools[20] was used to delineate the A and B compartments and to obtain the PC1 scores on 10kb binned matrices, 40kb binned matrices. The 10kb compartmentalization was used to initialize the chromatin polymer and the 40kb was used for compartment switching analysis. The Hi-C contact maps hardly show any differences between the control and TSA treated cells as shown in Fig. S18, S19. In Fig S20, we plot the compartment changes over other chromosomes chr18-21 to show that the compartment changes are a boundary phenomenon.

Calculating the flipping score from the simulations and matching with simulations: The flipping score was calculated by averaging scores across simulations. Initially, each chromatin segment is assigned a score of 0. For each simulation, segments changing to blue are assigned +1, while those changing to red are assigned -1. The scores are then summed across multiple simulations and averaged. The final flipping score represents the frequency of mark changes for each segment. An absolute threshold of 0.5 is applied to determine significance, indicating a segment has changed its mark in at least half of the simulations. Finally, the scores are compared to experimental results to assess their correlation.

##### 4.2 RNA-seq analysis for A375 cells

The RNA-seq data for the three replicates was combined and analyzed using DESeq2 followed by GO enrichment analysis from cluster profiler to get the differentially regulated pathways. The differential pathway analysis was repeated in the following two settings:

1. For all the genes (plotted in the main text)
2. For genes within 300kb of the domain boundaries (plotted in Fig. S21)

Additionally, we have also plotted the differentially expressed genes on chr19 in Fig. S22 which shows that differentially regulated genes are close to domain boundaries.

Lastly, we found that Wnt signaling pathway (GO:0060070) and positive regulation of stress-activated MAPK cascade (GO:0051403) were both enriched within 300kb more than outside of it. The gene lists from Table S2 provide the support for the percentages we plot in the main text.

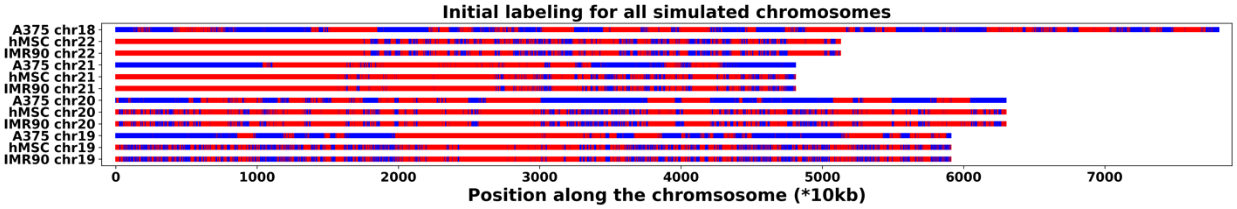


Fig. S1: Initial labeling of all simulated chromosomes.

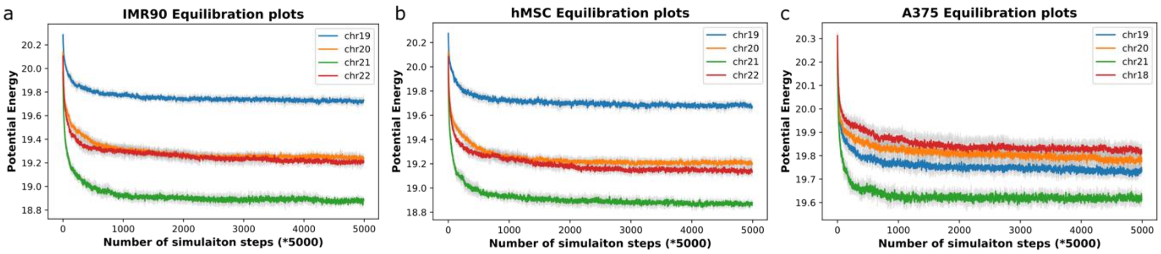


Fig. S2: Equilibration plots for all the simulated chromosomes.

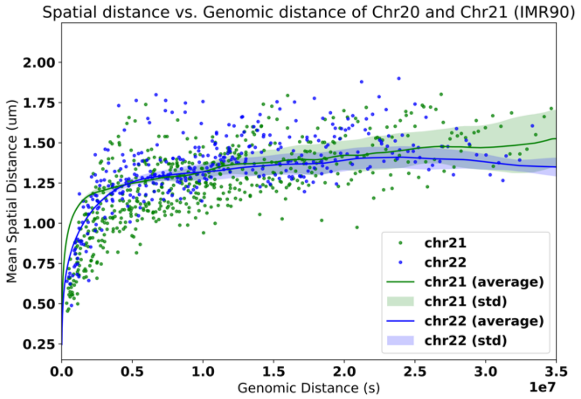


Fig. S3: Experimental validation of the chosen potentials. The scatter represents the experimentally observed values[1] while the lines and the shaded regions represents the simulated fit.

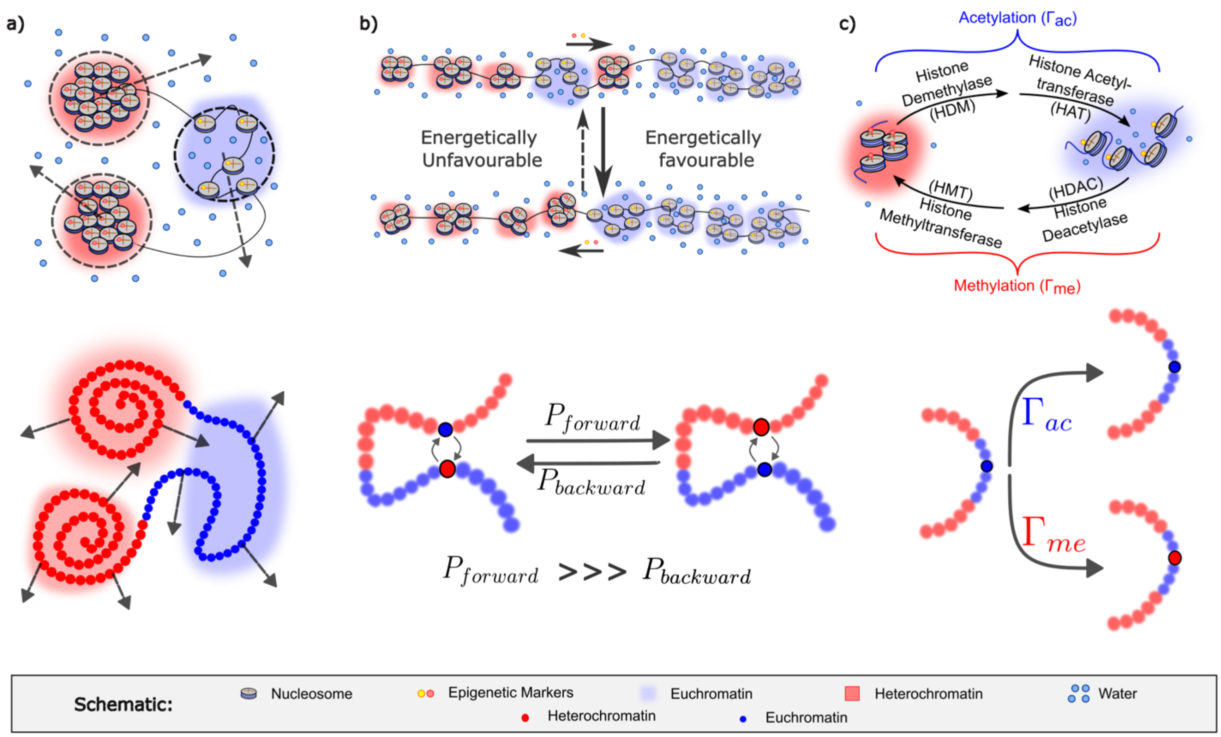


Fig. S4: Executed dynamics and corresponding segment-scale polymer representations of the dynamics with a) representing Brownian motion, b) representing diffusion of epigenetic marks and c) showing execulation of epigenetic reactions

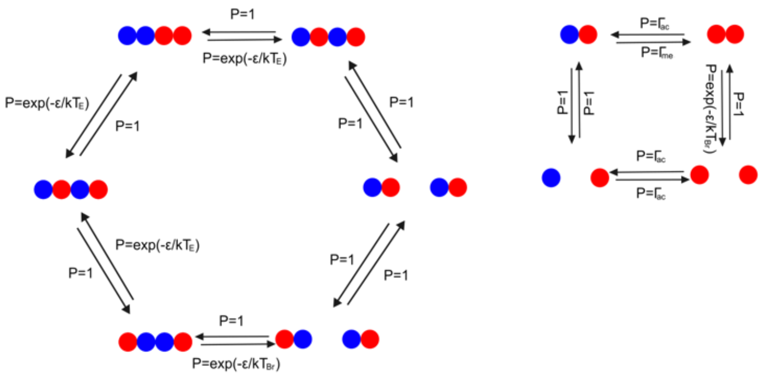


Fig. S5: Diffusion processes maintain detailed balance and reactions introduce out-of-equilibrium phenomenon.

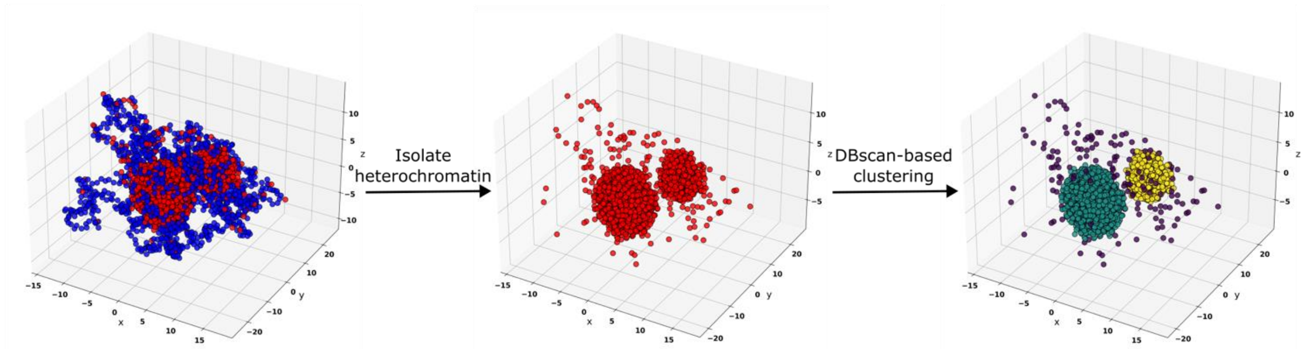


Fig. S6: Computing the radius of gyration of the domains involves isolating the heterochromatin regions followed by clustering using DBSCAN.

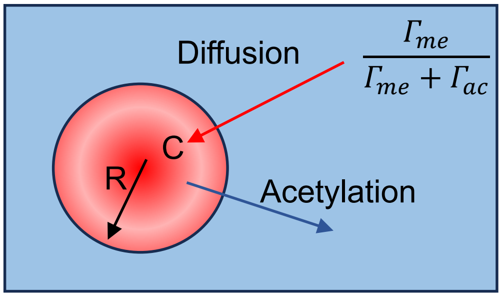


Fig. S7: Representative diffusion-reaction set-up at steady state

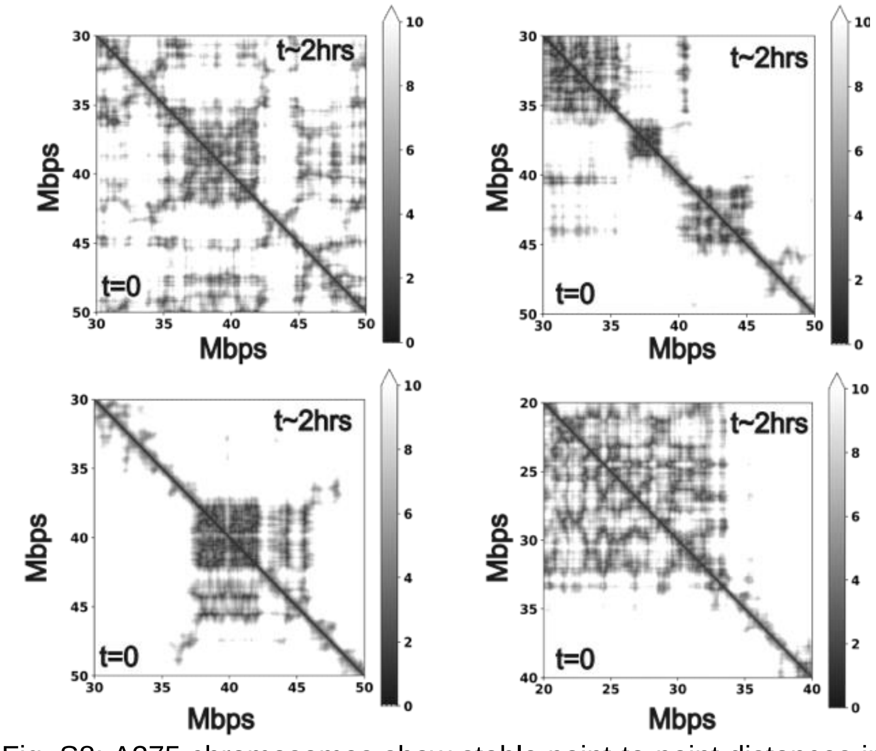


Fig. S8: A375 chromosomes show stable point-to-point distances in 2 hours of real time. Chr18, 19, 21 and 20 are placed clockwise. The scale is 100nm.

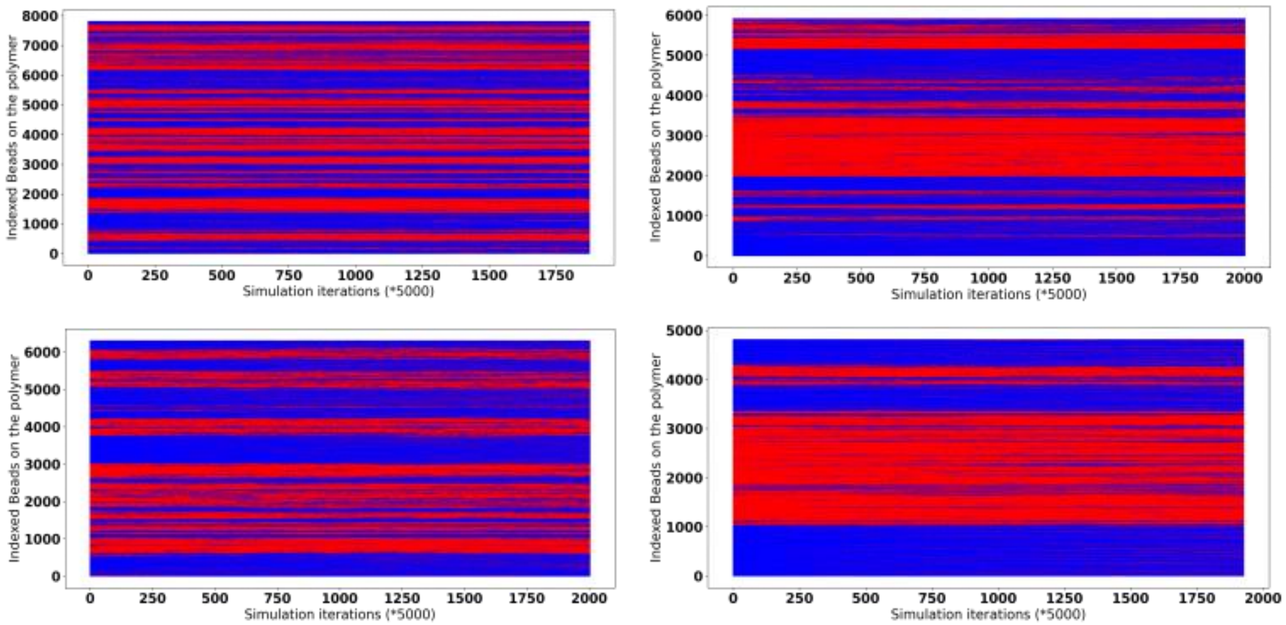


Fig. S9: Domain stability over 10 hours of simulation for A375 chromosomes. Chr 18, 19, 21 and 22 are placed clockwise

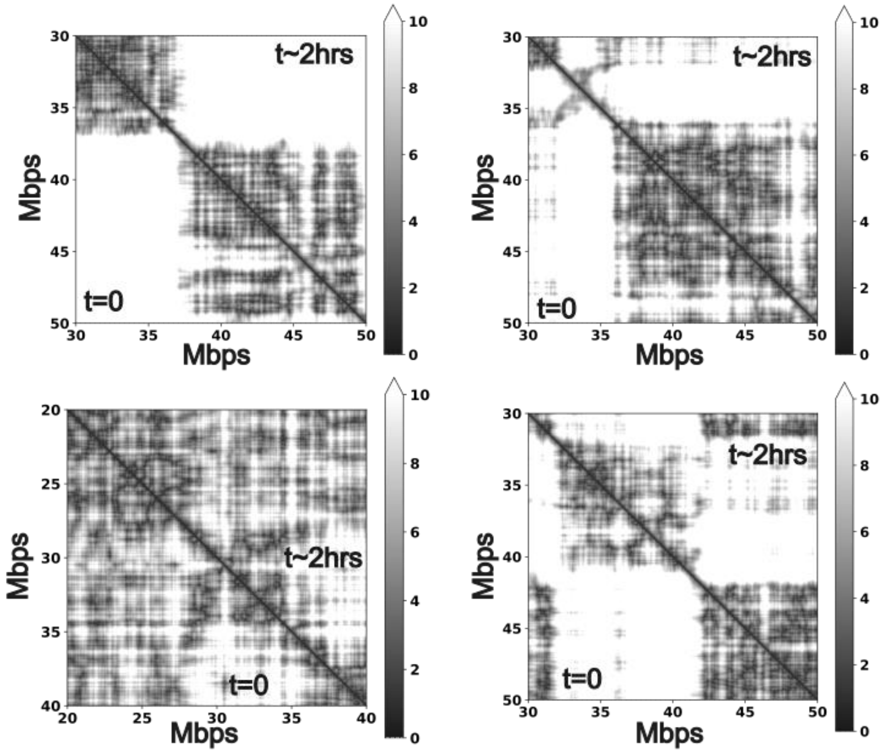


Fig. S10: hMSC chromosomes show stable point-to-point distances in 2 hours of real time. Chr19, 20, 22 and 21 are placed clockwise. The scale is 100nm.

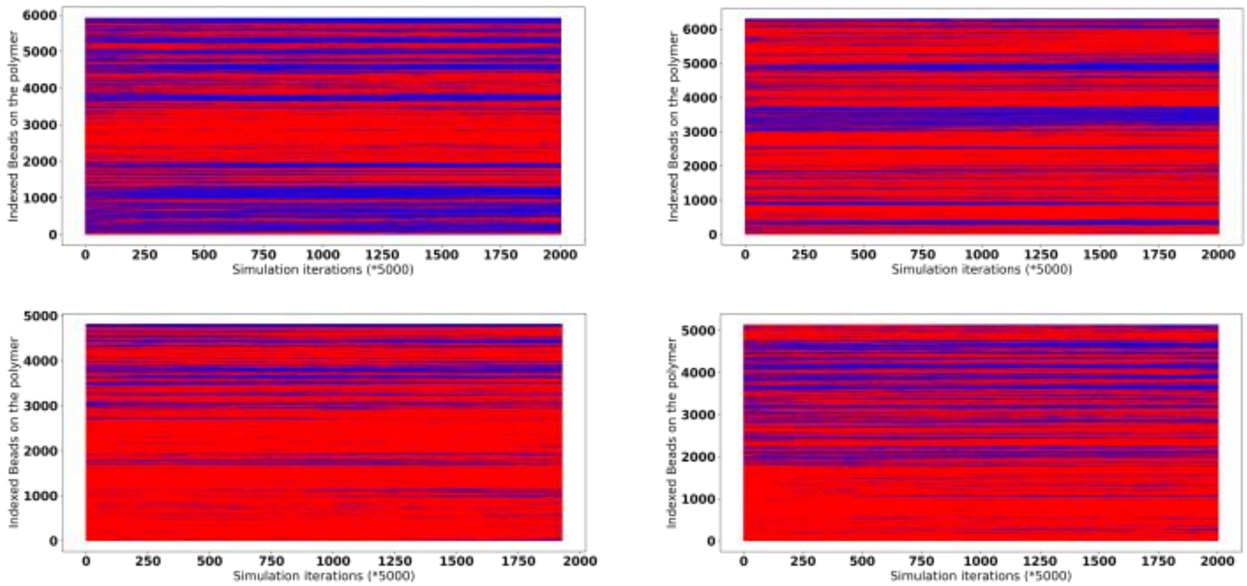


Fig. S11: Domain stability over 10 hours of simulation for hMSC chromosomes. Chr 19, 20, 22 and 21 are places clockwise.

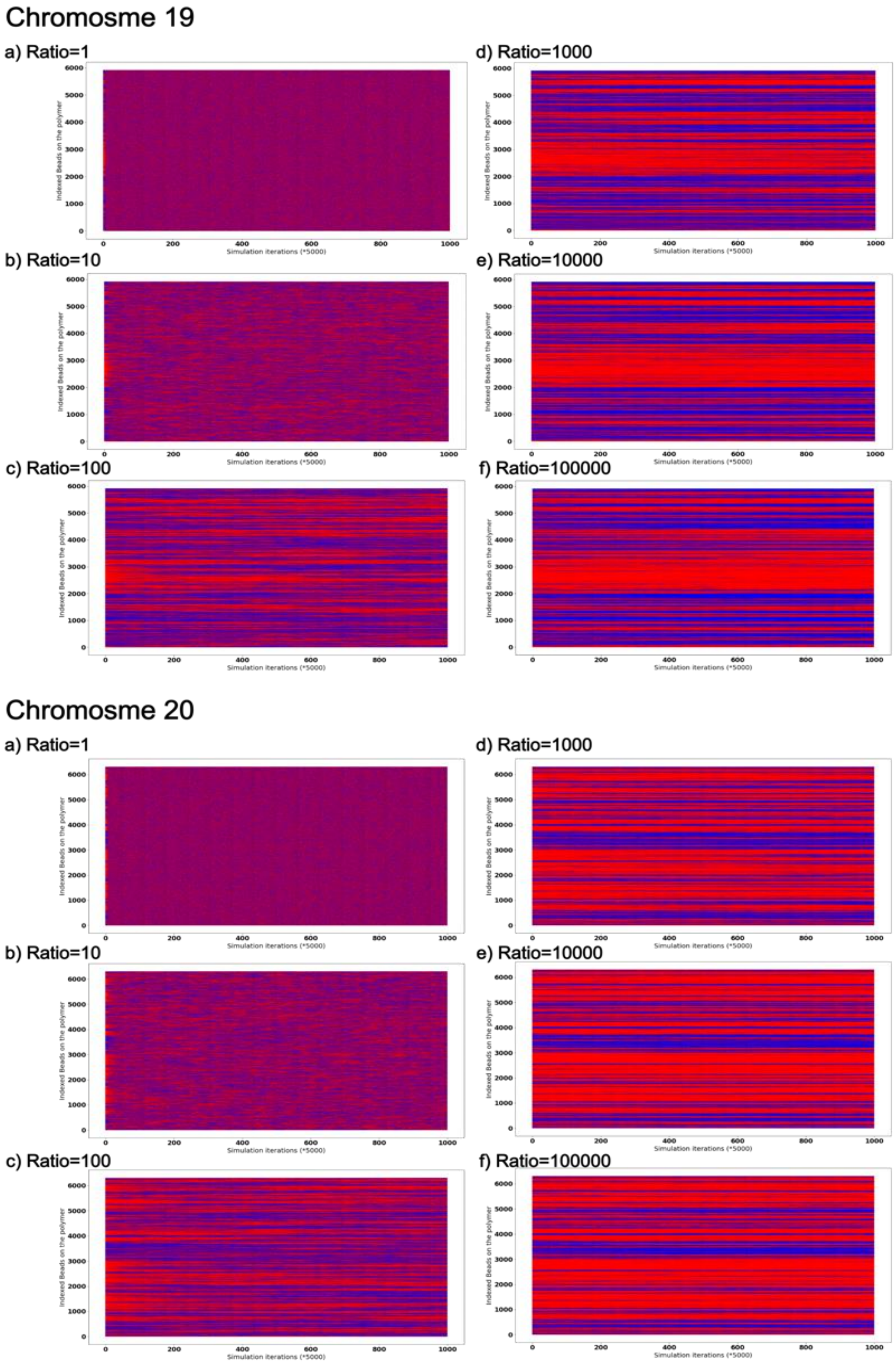


Fig. S12: Kymographs of evolution of the chromatin domains on the linear polymer for chromosomes 19 and 20 of hMSC. The plots show the evolution of the domains with time at different ratios of number of steps at which epigenetic reaction takes place to the number of steps at which epigenetic diffusion takes place.

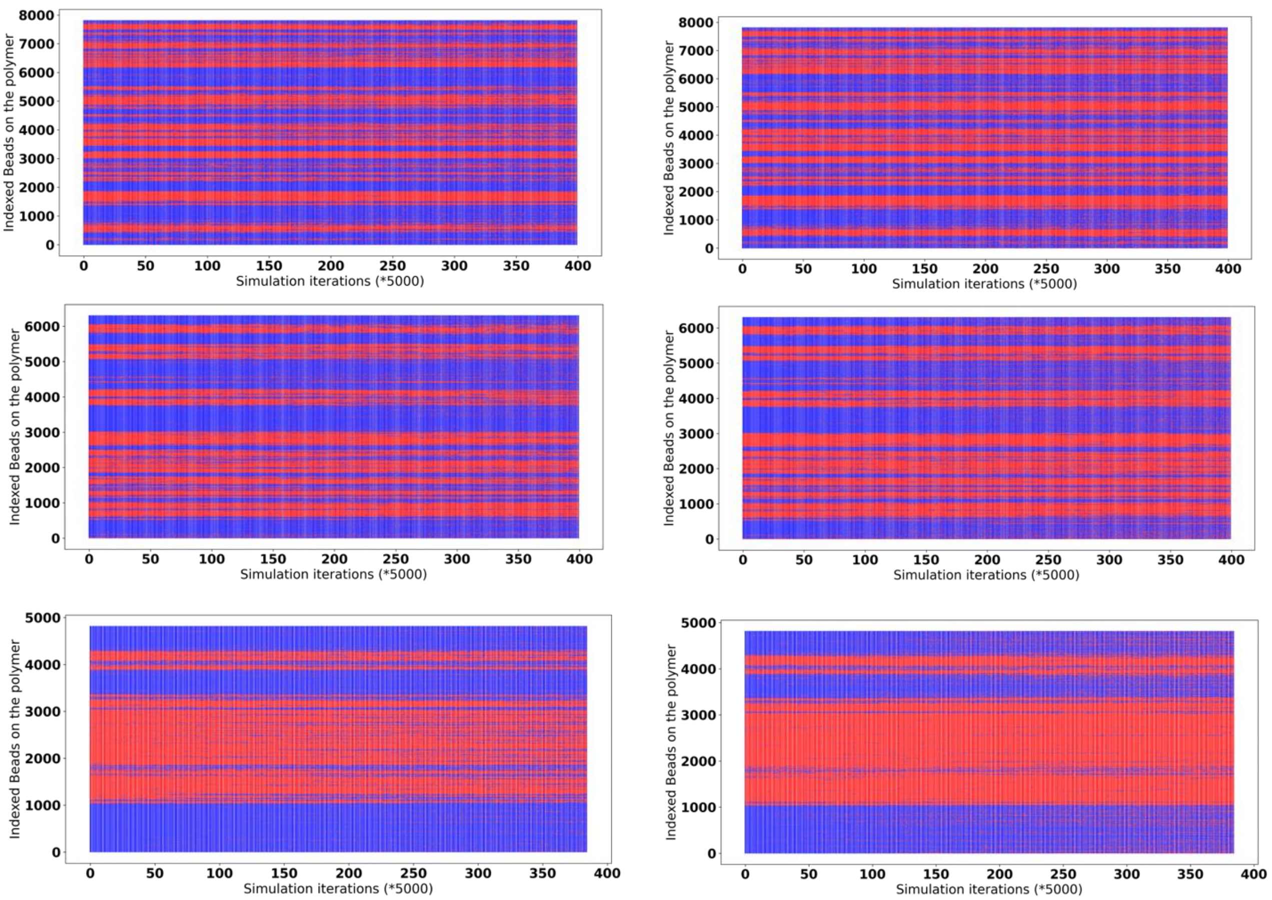


Fig. S13: Kymographs of evolution of the epigenetic landscape for A375 chromosomes. Each row is a single chromosome with higher acetylation ($\Gamma_{me}/\Gamma_{ac}$ =0.2) in the left panel and higher methylation ($\Gamma_{me}/\Gamma_{ac}$ =0.9) in the right panel. From top to bottom: Chr18, Chr20 and Chr22.

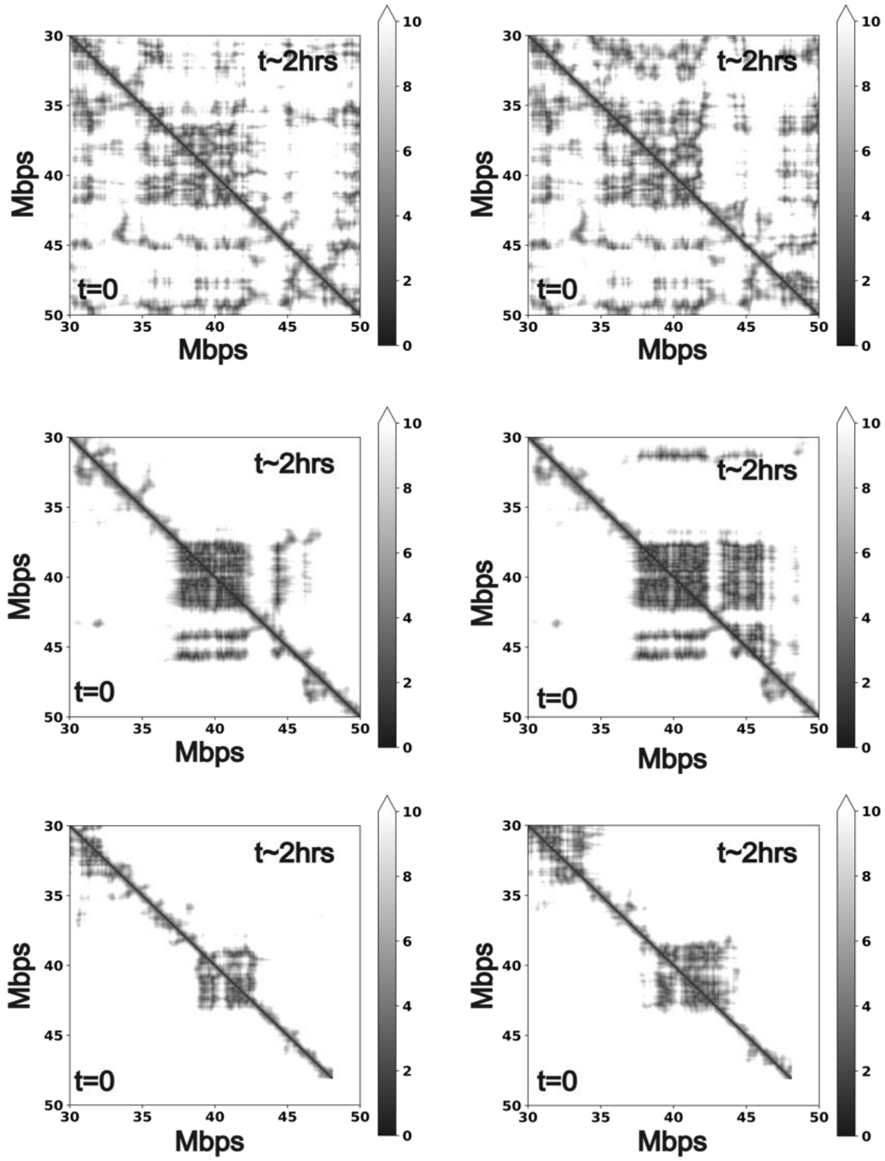


Fig. S14: A375 distance maps for acetylation and methylation. Each row is a single chromosome with the left panel showing acetylation ($\Gamma_{me}/\Gamma_{ac}$ =0.2) and the right panel showing methylation($\Gamma_{me}/\Gamma_{ac}$ =0.9). From top to bottom: Chr18, Chr20, Chr21. The scale is 100nm.

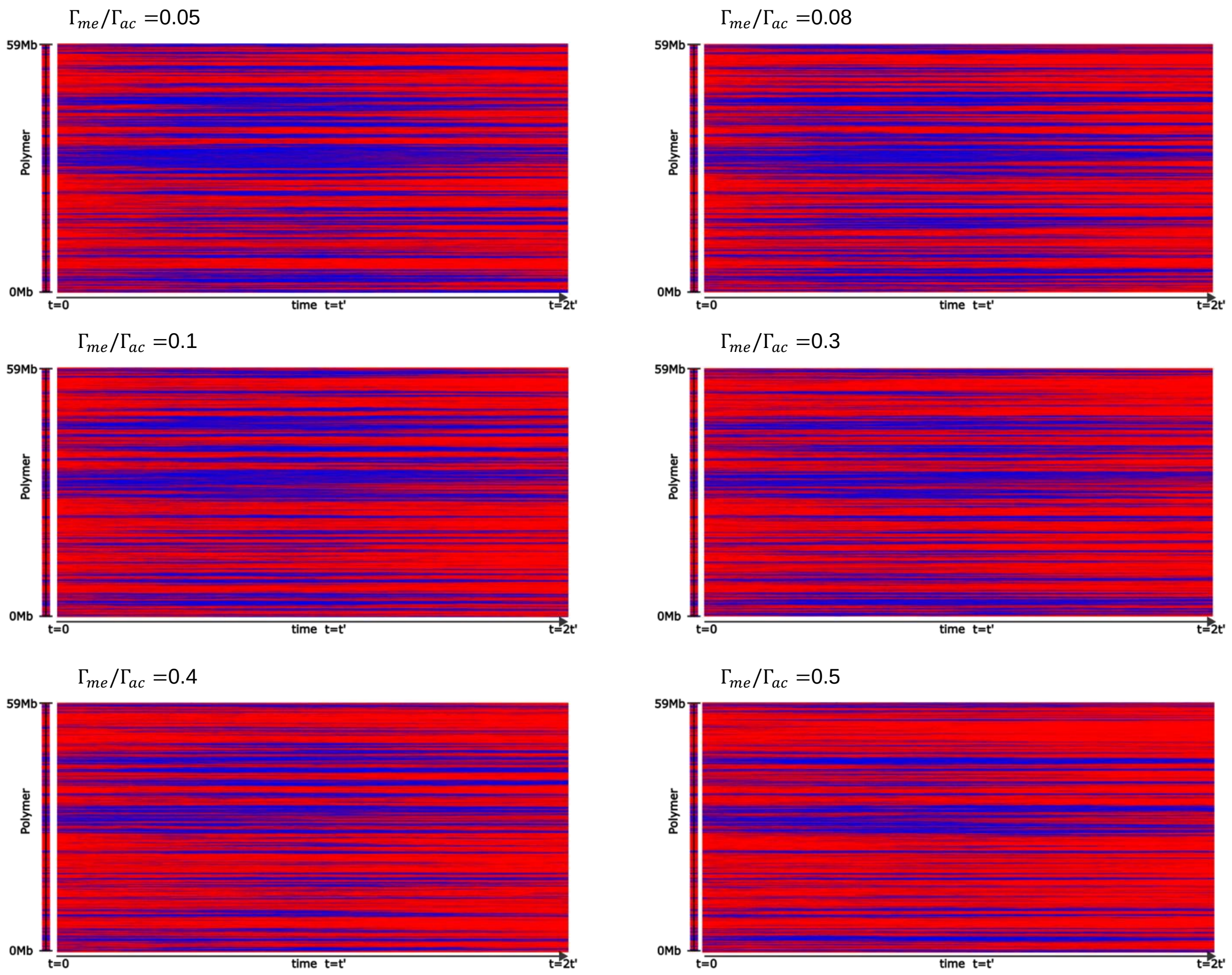


Fig. S15: Memory runs for hMSC Chr20. For all runs the initial and final reaction ratios are the same to represent the initial conditions ($\Gamma_{me}/\Gamma_{ac}$). At the intermediate point the ratio is ($\Gamma_{me}/\Gamma_{ac}$) is mentioned with the plot.

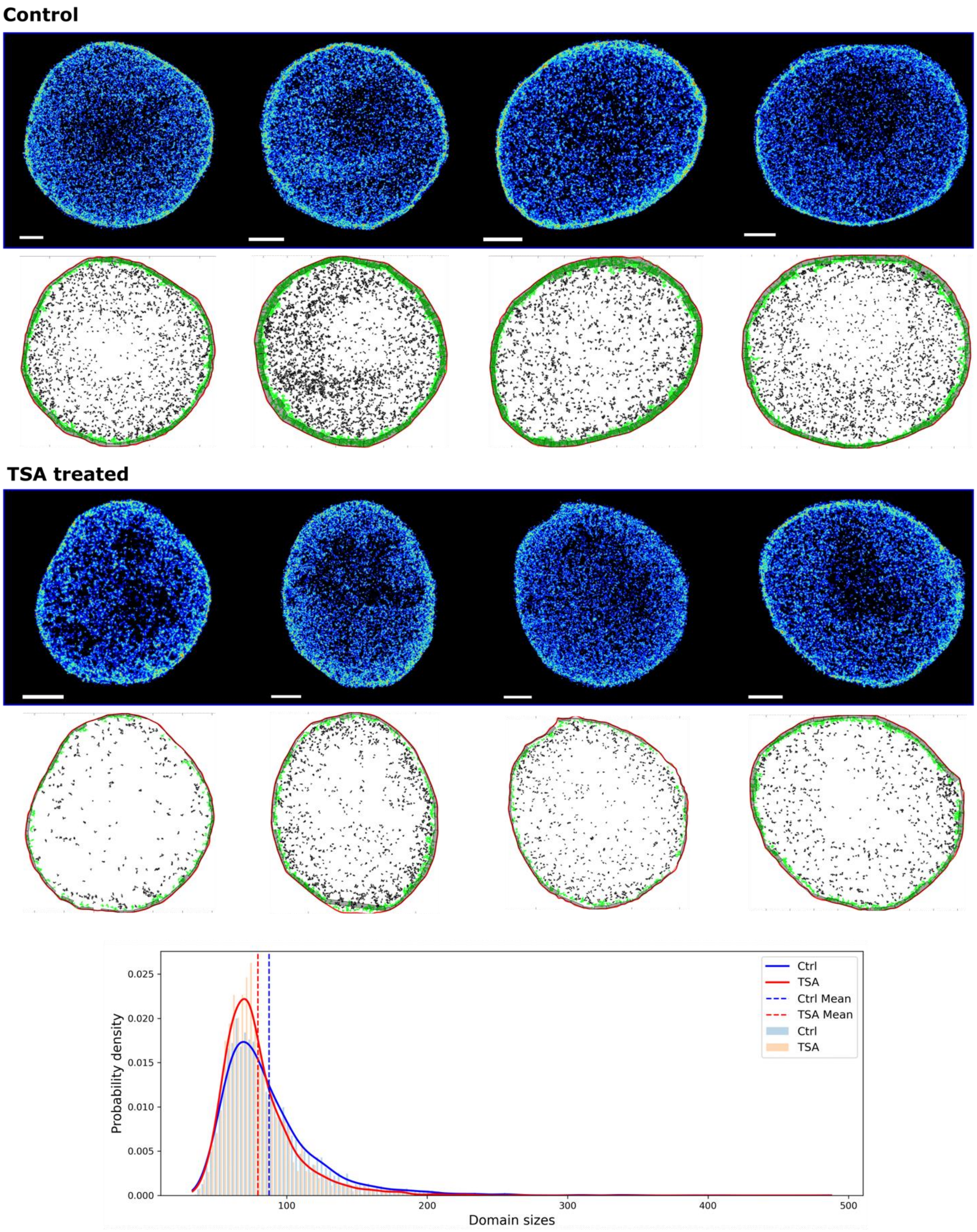


Fig. S16: STORM analysis for control and TSA treated A375 cells. For each cell, shown in the black panel, its corresponding DBSCAN detected interior domains are shown as black clusters. The lamin associated domains (in green boxes) are excluded from the analysis. The final size distributions for the shown images are provided in the bottom panel along with the means.

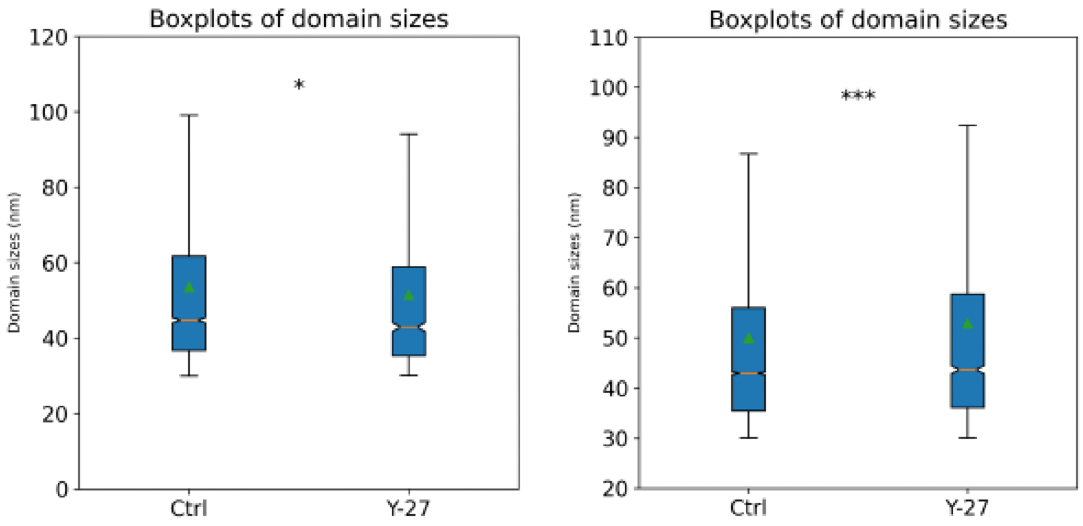


Fig. S17: Chromatin domain sizes scale with Y-27 treatment in hMSCs and are affected in tendinosis[1].

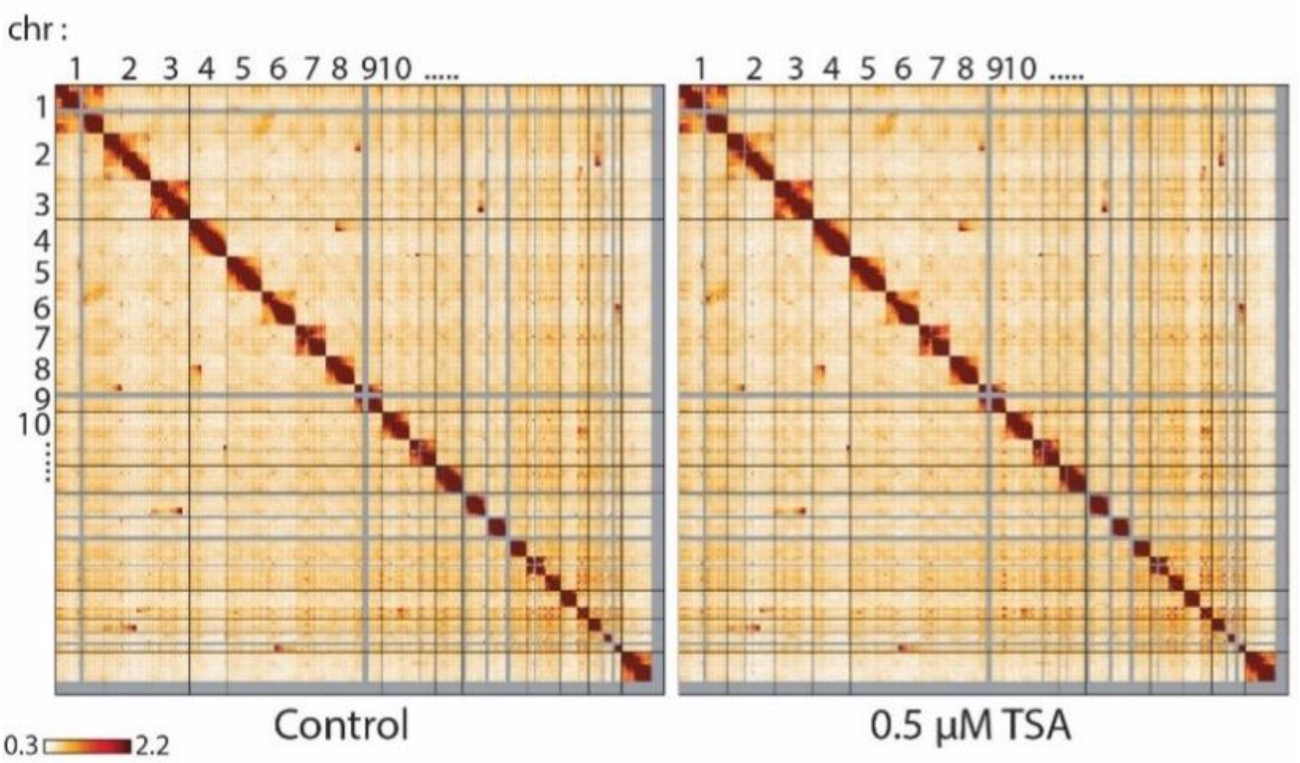


Fig. S18: Control and TSA treated Hi-C matrices for all chromosomes.

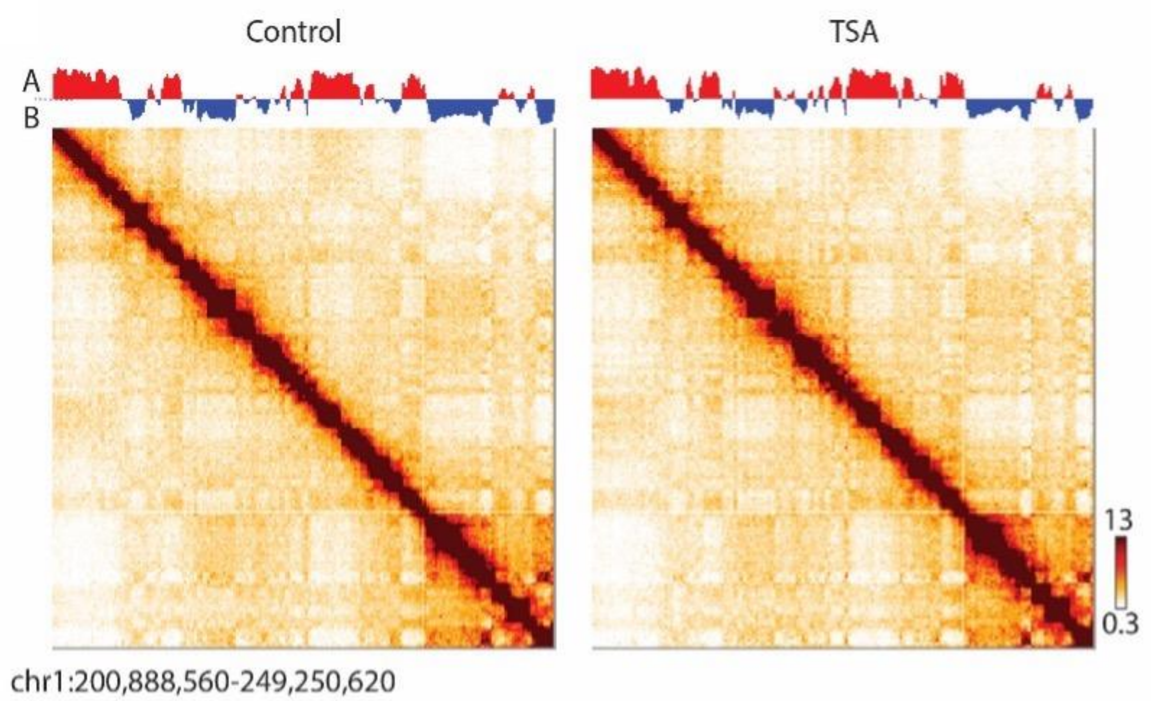


Fig. S19: HiC maps do not show any visible changes between control and TSA treated cells for A375 cells.

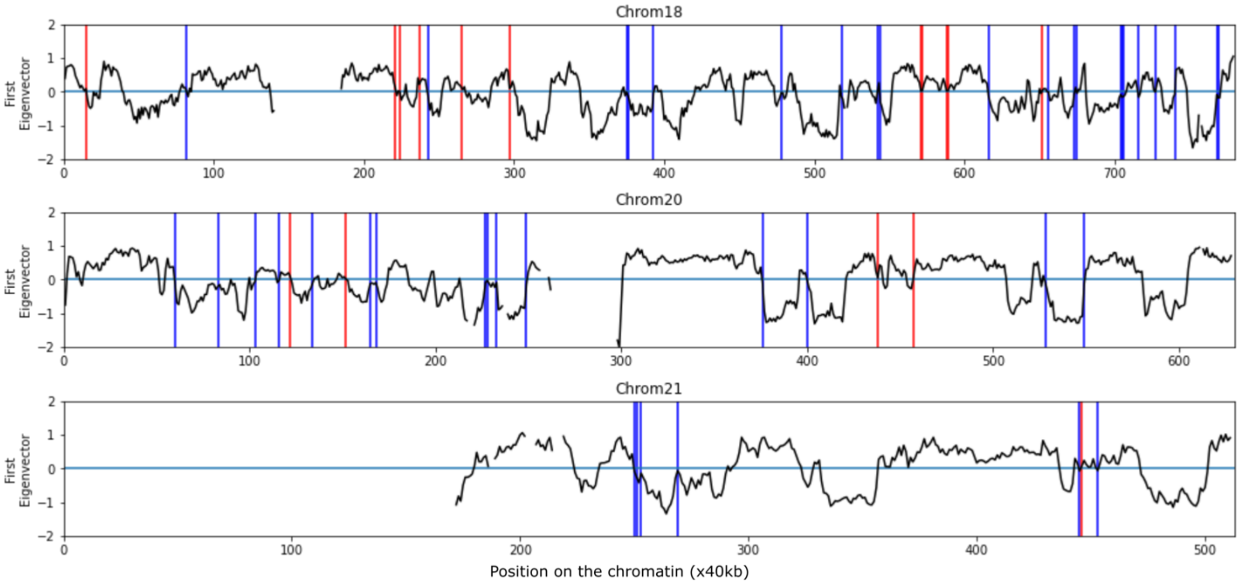


Fig. S20: Compartment switches for chr18, 20 and 21 for A375 cells after TSA treatment. The black lines represent the control PC1 with the blue vertical lines representing B to A transition and red vertical lines representing A to B transitions.

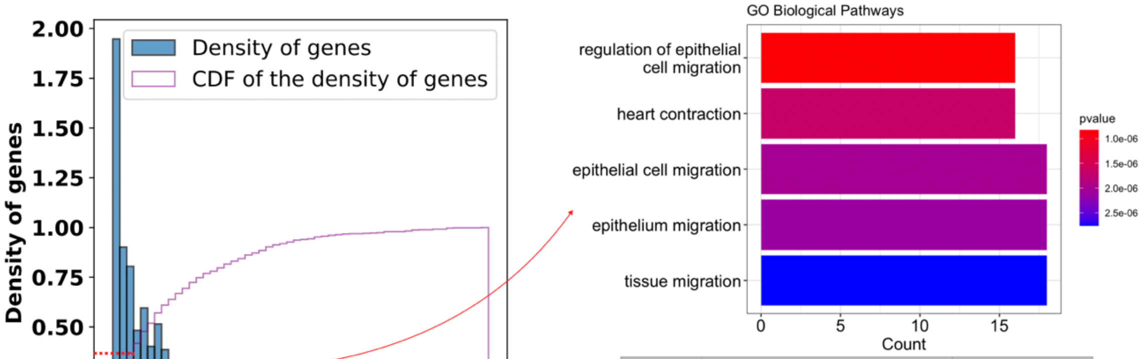


| ID | Description | pvalue |
|---|---|---|
| GO:0010634 | positive regulation of epithelial cell migration | 3.33147509029984E-05 |
| GO:0043410 | positive regulation of MAPK cascade | 5.26583341551882E-05 |
| GO:0060070 | canonical Wnt signaling pathway | 5.74718866581155E-05 |
| GO:0016055 | Wnt signaling pathway | 2.30133348572586E-04 |
| GO:0070371 | ERK1 and ERK2 cascade | 2.7428406078365E-04 |

Fig. S21: Almost 30% differentially regulated genes lie within 300kbps of the domain boundaries. GO analysis of these genes using clusterProfiler shows that migration and ERK, Wnt pathways are still majorly upregulated.

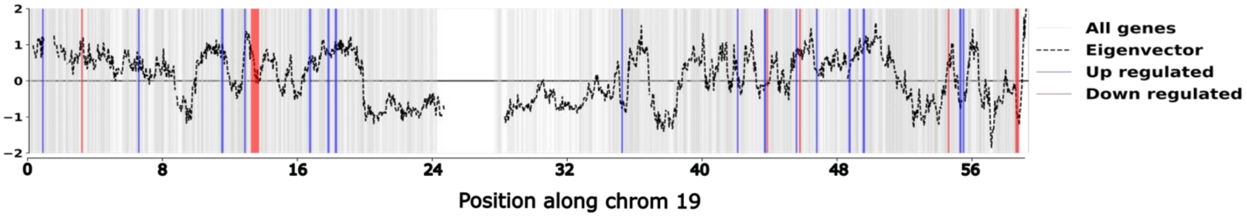


Fig. S22: Significantly differentially regulated genes are placed close to the domain boundaries.

| *Cell line* | Source | Chromosomes | Purpose |
|---|---|---|---|
| *IMR90* | Epigenomics Roadmap ChromHMM | 19,20,21 | To realize the correct potentials |
| *A375* | Experimentally obtained Hi-C contact maps | 18,19,20,21 | To match with STORM imaging and Hi-C changes upon TSA treatment |
| *hMSC* | Epigenomics Roadmap ChromHMM | 19,20,21,22 | To study the effect of substrate stiffness change and GSK treatment |

Table S1: Data for specific cell lines and their usage

| **Pathway** | **Genes within 300kb** | **Ratio of genes found of this pathway found in 300kb /total genes within 300kb** | **Total number of genes for this pathway/ total number of genes** |
|---|---|---|---|
| **Wnt signaling pathway** | FZD4/ TMEM170B/ GSKIP/ STK4/ WNT7B/ CCNYL1/ SOX9/ SCYL2/ EXT1/ CYLD/ LGR4/ CTNND1/ PRICKLE1/ WNT2B | 14/227 | 1.6% |
| **positive regulation of stress-activated MAPK** | TRAF1/ PER1/ MAPK8IP1/ MAP2K3/ TNIK/ MAP3K3/ TLR9/ | 7/227 | 1.5% |

Table S2: MAPK and Wnt signaling pathway enrichment within 300kb
